# The conserved histone chaperone Spt6 facilitates DNA replication and mediates genome instability

**DOI:** 10.1101/2022.04.27.489770

**Authors:** Catherine LW Miller, Fred Winston

## Abstract

Histone chaperones are an important class of proteins that regulate chromatin accessibility for DNA-templated processes. Spt6 is a conserved histone chaperone and key regulator of transcription and chromatin structure. However, its functions outside of these roles have been little explored. In this work, we demonstrate a role for *S. cerevisiae* Spt6 in DNA replication and more broadly as a regulator of genome stability. Spt6 binds the replication machinery and depletion or mutation of Spt6 impairs DNA replication *in vivo*. Additionally, *spt6* mutants are sensitive to DNA replication stress inducing agents, with increased sensitivity when combined with loss of DNA replication associated factors. Furthermore, *spt6* mutants have elevated levels of DNA double strand breaks and recombination. These effects appear to be independent of R-loops, which are not elevated in *spt6* mutants. Our results identify Spt6 as a regulator of genome stability, at least in part through a role in DNA replication.

## INTRODUCTION

Histone chaperones are conserved eukaryotic factors that directly regulate histone-DNA interactions to enable transcription, DNA replication, and DNA repair (Hammond et al., 2017; Warren and Shechter, 2017). They function in cooperation with histone modifying enzymes and chromatin remodelers to coordinate the dynamic opening of chromatin to allow DNA-templated activities, while also modulating the closing of chromatin to prevent the loss of genome integrity and to maintain epigenetic memory. Insights into the functions of histone chaperones are important to understand chromatin-regulated processes as well as how mutations that alter histones and histone chaperones contribute to cancer and other human diseases (Nacev et al., 2019; Plass et al., 2013; Ray-Gallet and Almouzni, 2022).

Spt6 is a highly conserved histone chaperone of great biological importance. It is essential for viability in *S. cerevisiae* (Clark-Adams and Winston, 1987; Neigeborn et al., 1987) and it is required for proper development in *C. elegans* (Nishiwaki et al., 1993), zebrafish (Keegan et al., 2002; Kok et al., 2007), and *Drosophila* (Ardehali et al., 2009; Bourbon et al., 2002). In mammalian systems, Spt6 is required for embryonic stem cell maintenance, muscle cell differentiation, epithelial cell differentiation, and immunoglobulin class-switch recombination (Begum et al., 2012; Okazaki et al., 2011; Vo et al., 2021; Wang et al., 2013, 2017). Additionally, loss-of-function alleles of the human homolog, *SUPT6H*, are greatly underrepresented (Karczewski et al., 2020), indicating it is likely essential in humans.

Spt6 is a large multi-domain protein that is a vital component of the transcription elongation complex (Close et al., 2011; Vos et al., 2018). The C-terminal tandem SH2 domains of Spt6 directly interact with phosphorylated residues in the carboxy-terminal domain (CTD) and linker region of Rpb1, the largest subunit of RNA Polymerase II (RNAPII) (Close et al., 2011; Diebold et al., 2010a; Sdano et al., 2017; Sun et al., 2010; Yoh et al., 2007). Furthermore, Spt6 directly interacts with histones and nucleosomes, and can assemble nucleosomes *in vitro* (Bortvin and Winston, 1996; McCullough et al., 2015; McDonald et al., 2010). As such, Spt6 plays critical roles regulating transcription initiation (Adkins and Tyler, 2006; Ivanovska et al., 2011), transcription elongation (Ardehali et al., 2009; Endoh et al., 2004; Narain et al., 2021; Žumer et al., 2021), repression of intragenic transcription (Cheung et al., 2008; Doris et al., 2018; Gouot et al., 2018; Kaplan et al., 2003; Uwimana et al., 2017), and transcription termination (Kaplan et al., 2005; Narain et al., 2021; Nojima et al., 2018). At transcribed genes, Spt6 is required to establish and/or maintain normal nucleosome positioning (van Bakel et al., 2013; Bortvin and Winston, 1996; DeGennaro et al., 2013; Doris et al., 2018; Ivanovska et al., 2011; Jeronimo et al., 2019; Perales et al., 2013) as well as for certain histone modifications, including methylation of H3K4 (Begum et al., 2012; DeGennaro et al., 2013; Kato et al., 2013; Petruk et al., 2006), H3K27 (Chen et al., 2012; Wang et al., 2013, 2017), and H3K36 (Carrozza et al., 2005; DeGennaro et al., 2013; Gopalakrishnan et al., 2019; Yoh et al., 2008; Youdell et al., 2008). Combined, these works demonstrate the important roles Spt6 plays in regulating eukaryotic transcription. However, the roles of Spt6 outside of transcription remain poorly understood.

Several lines of evidence have hinted that Spt6 may also have a role in DNA replication. Yeast *spt6* mutants have phenotypes consistent with DNA replication defects, such as an increase in recombination frequency (Malagon and Aguilera, 1996; Malagón and Aguilera, 2001), chromosome segregation errors (Basrai et al., 1996), and sensitivity to the DNA replication inhibitor hydroxyurea (HU) (Diebold et al., 2010b; Viktorovskaya et al., 2021). Consistent with this potential role, depletion of Spt6 from mammalian cells decreases total DNA synthesis (Narain et al., 2021). However, no studies have been performed to directly test the possibility that Spt6 is required for DNA replication.

In this study, we provide evidence that Spt6 has a direct role in DNA replication. First, DNA replication is impaired *in vivo* either after Spt6 depletion or in *spt6* mutants. The role of Spt6 in replication is likely direct, as Spt6 physically interacts with the DNA replication helicase, dependent on the SH2 domains of Spt6. In addition, *spt6* mutants are synthetically sick with loss of the DNA replication factor Ctf4, as well as with DNA replication checkpoint mutants. Finally, we suggest that RNA-DNA hybrids are not driving genome instability in *spt6* mutants, as we do not observe a correlation between RNA-DNA hybrid levels and genome instability. Our work underscores the importance of Spt6 in mediating genome stability and identifies a role for Spt6 in DNA replication in addition to its well-established role in transcription.

## RESULTS

### Spt6 Depletion Causes DNA Replication Defects

To examine the requirement for Spt6 in DNA replication, we first tested whether loss of Spt6 impaired DNA replication *in vivo*. As Spt6 is essential for viability in *S. cerevisiae*, yeast strains were constructed in which we could deplete Spt6 using an auxin-inducible degron (AID) system (Doris et al., 2018; Nishimura et al., 2009) and then isolate newly synthesized DNA by measuring the incorporation of the thymidine analog BrdU (Lengronne et al., 2001; Sivakumar et al., 2004). We synchronized cells in G1 with the yeast mating pheromone alpha factor, followed by treatment with the auxin indole-acetic acid (IAA) to deplete Spt6-AID, or with DMSO as a non-depletion control (Figure 1A). In the depleted samples, Spt6 protein levels were reduced to ∼10% of non-depleted levels and cells remained arrested in G1 following depletion (Figure S1A, B). Importantly, cell viability was greater than 88% in all Spt6-depleted samples. After release from alpha factor arrest, samples were harvested at an early time point in S phase (30 minutes). Newly replicated DNA was isolated by immunoprecipitation with an anti-BrdU antibody followed by qPCR at well-characterized early and late yeast replication origins (ARSs). To allow for normalization between samples, equal amounts of BrdU-labeled *S. pombe* genomic DNA were spiked into each sample prior to immunoprecipitation.

**Figure 1.**
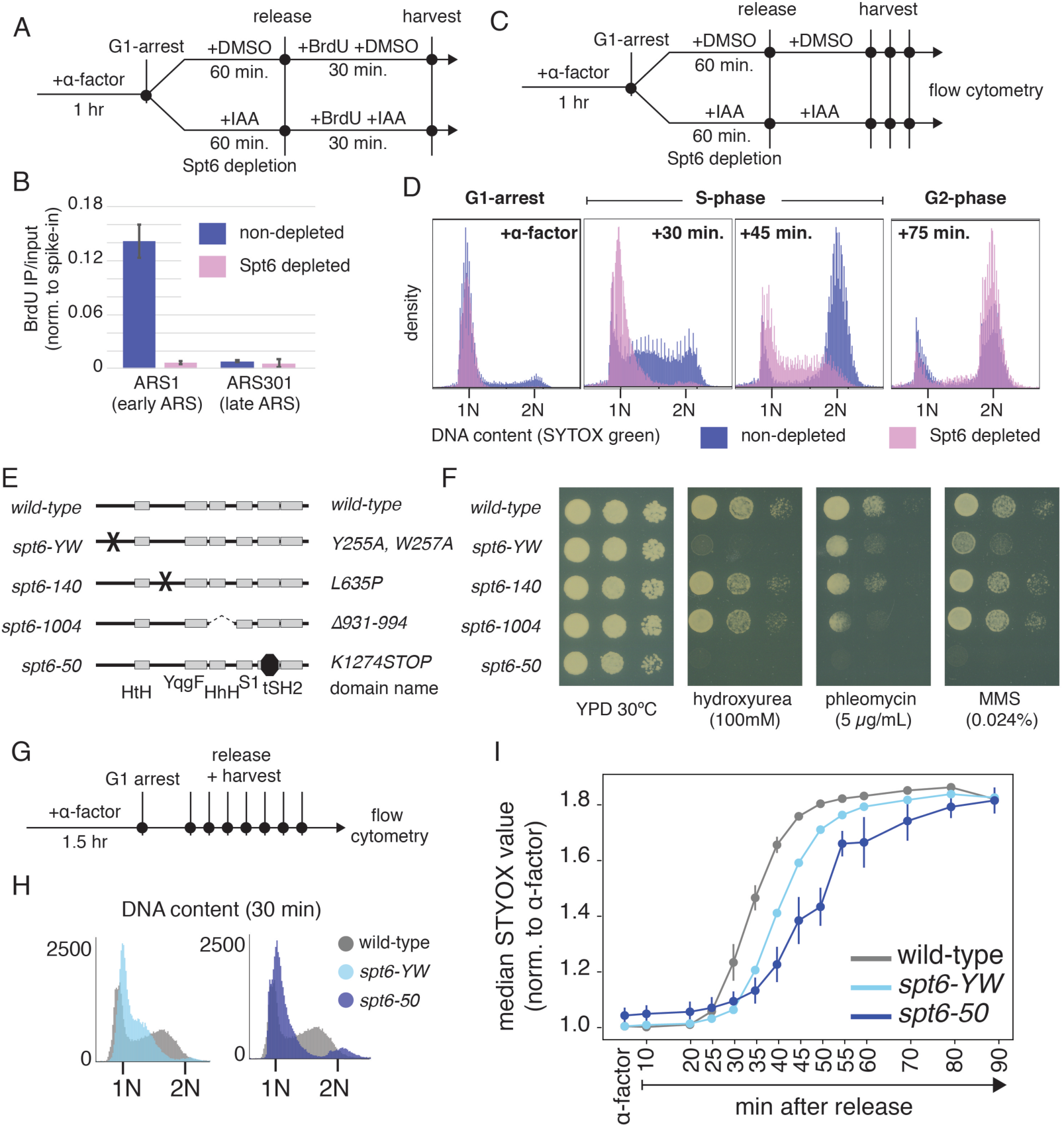
Spt6 depletion and *spt6* mutants have DNA replication defects. (A) Schematic of cell synchronization, Spt6 depletion, and harvest strategy for the experiment shown in panel B. (B) BrdU-IP-qPCR IP/input (n=2), data are normalized to 10% BrdU-labeled *S. pombe* genomic DNA. (C) Schematic of cell synchronization, Spt6 depletion, and harvest strategy for the experiment shown in panel D. (D) Samples were stained with SYTOX green and total DNA content was analyzed by flow cytometry. (E) Schematic of *spt6* mutations. The conserved domains (Close et al., 2011) are indicated as gray boxes and labeled at the bottom. An “X” indicates a point mutation, a dashed line indicates a deleted region, and an octagon indicates a nonsense mutation. The amino acid changes caused by each mutation are indicated to the right of the diagram. (F) Serial dilution plating of *spt6* mutants. Strains were serially diluted 10-fold, spotted onto the indicated plates, and incubated at 30°C for three days. (G) Schematic of cell synchronization, Spt6 depletion, and harvest strategy for the experiments shown in G and H. (H) Flow cytometry analysis was performed for wild-type (n=2), *spt6-YW* (n=2) and *spt6-50* (n=3). The median fluorescence signal over time was plotted, normalized for the signal in alpha-factor arrested cells. (I) Representative flow cytometry histogram of total DNA content for the indicated strains at 30 minutes following release from alpha-factor. The median values with standard deviation were plotted vs. time.

Our results showed a requirement for Spt6 for DNA synthesis. In the non-depleted samples there was a high level of BrdU incorporation at the early replication origin *ARS1*, but not at the late-firing origin *ARS301,* as expected. In contrast, in the Spt6-depleted samples, BrdU incorporation was undetectable at either origin (Figure 1B). These results are in agreement with recent work in mammalian cells where depletion of Spt6 also resulted in loss of BrdU incorporation (Narain et al., 2021).

To test whether the Spt6-depleted cells had arrested growth rather than impaired DNA synthesis, we also assayed S-phase progression following Spt6 depletion by monitoring total DNA content by flow cytometry (Figure 1C). Our results showed that Spt6-depeleted cells completed S phase but with a delay in DNA synthesis. The median fluorescence value for non-depleted samples at 30 minutes was similar to the median fluorescence value for Spt6-depleted cells at 45 minutes. By 75 minutes, the Spt6-depleted cells had completed DNA synthesis, showing that cells are capable of completing at least one cell cycle in the absence of Spt6 (Figure 1D). These data indicate that Spt6 has a role in DNA replication.

### *spt6* Mutants Are Sensitive to Chemicals That Cause DNA Replication Stress

To investigate the requirement for Spt6 in DNA replication in greater depth, we analyzed *spt6* mutants. The *spt6* mutants allow for continuous cell division at 30°C, in contrast to Spt6 depletion, which eventually leads to inviability. For these experiments, we chose four *spt6* mutations (Figure 1E): (i) *spt6-YW* (Diebold et al., 2010b; Viktorovskaya et al., 2021), (ii) *spt6-140* (Malagón and Aguilera, 2001; Winston et al., 1984) (iii) *spt6-1004* (Doris et al., 2018; Kaplan et al., 2003), and (iv) *spt6-50* (Diebold et al., 2010b). Previous studies showed that these mutations cause distinct phenotypes with respect to effects on transcription, histone modifications, and nucleosome organization (Chu et al., 2006; Doris et al., 2018; Dronamraju and Strahl, 2014; Viktorovskaya et al., 2021; Youdell et al., 2008). For each allele, the mutant proteins are present at near wild-type levels at 30°C (Figure S1C).

We note that our previous analysis of the *spt6-YW* mutant showed that, during growth at 30°C, there was little change in transcript levels, including genes encoding the G1 cyclins or ribonucleotide reductase (Figure S1D, E) (Viktorovskaya *et al*., 2021, data available at GEO:GSE160821). Therefore, for the *spt6-YW* mutant, any defects in DNA replication are unlikely to be the consequence of transcriptional changes.

As a first step to understand if these *spt6* mutants have defects in DNA replication, we tested their sensitivity to agents that induce DNA replication stress, including hydroxyurea, phleomycin, and methylmethanesulfonate. Our results showed allele-specific sensitivities to these agents, with *spt6-YW* and *spt6-50* being the most sensitive (Figure 1F, Figure S3), in agreement with previous reports (Dronamraju and Strahl, 2014; Viktorovskaya et al., 2021). We chose to proceed with further analysis of the *spt6-YW* and *spt6-50* alleles.

### *spt6* Mutants Have S-phase Progression Defects

As we had observed a delayed S-phase progression after Spt6 depletion, we asked whether this delay was also present in *spt6* mutants. Similar to our previous experiments, we synchronized cells in G1, released them into fresh medium, and harvested samples for analysis of total DNA content by flow cytometry (Figure 1G). Samples were taken every 10 minutes throughout one 90-minute cell cycle, increasing to 5-minute intervals during S phase (30-60 minutes). From these results, we observed an increase in DNA content over time in wild-type cells that follows the well-defined timing of the yeast cell cycle. However, both the *spt6-YW* and *spt6-50* mutants exhibited a delay in DNA synthesis compared to wild-type, apparent from the shift in curve of median fluorescent intensity values (Figure 1H). This delay is also apparent by plotting the distribution at 30 minutes, early S phase, where we observed an increased number of cells between 1N and 2N content for both mutants, similar to what we observed after Spt6 depletion (Figure 1I).

Combined, we observed that loss or impairment of Spt6 function causes DNA replication defects. To better characterize these defects, we focused on the *spt6-YW* and *spt6-50* mutants.

### *spt6* Mutants Have DNA Replication Initiation Defects *in vivo*

We hypothesized that DNA replication could be impaired by several different molecular mechanisms. Among these possibilities, we considered a requirement for Spt6 for a normal level of initiation or elongation by DNA polymerases, similar to how Spt6 regulates initiation and elongation of RNA polymerases (Adkins and Tyler, 2006; Ardehali et al., 2009; Doris et al., 2018; Endoh et al., 2004; Ivanovska et al., 2011; Narain et al., 2021; Žumer et al., 2021). In this general model, the impairment in *spt6* mutants would decrease the level of DNA synthesis around known origins. We also considered the possibility that Spt6 controls the fidelity of DNA replication initiation, similar to how Spt6 prevents intragenic initiation of transcription from within gene bodies (Cheung et al., 2008; DeGennaro et al., 2013; Doris et al., 2018; Gouot et al., 2018; Kaplan et al., 2003; Uwimana et al., 2017; Viktorovskaya et al., 2021). By this model, Spt6 impairment would result in initiation of replication from ectopic or dormant sites, thereby diluting replication factors and reducing the overall level of functional replication.

To differentiate between these possibilities, we measured the level and genomic location of newly synthesized DNA by measuring BrdU incorporation in wild-type and *spt6* mutant strains. Cells were first synchronized in G1 by treatment with alpha factor, and then released into medium containing 0.2 M HU and BrdU for one hour, resulting in arrest in early S phase (Figure 2A). After the HU arrest, DNA was extracted, BrdU-labeled DNA was immunoprecipitated, and next-generation sequencing libraries were prepared. For normalization between libraries, we used a spike-in control of BrdU-labeled *S. pombe* DNA. Biological duplicates were performed for each strain and the replicates correlated well, with Pearson correlations for spike-in normalized IP signal at all loci ranging from 0.93 – 0.99 (Figure S2A). Importantly, we showed that the one-hour treatment with 0.2 M HU did not greatly affect *spt6* mutant cell viability despite the HU sensitivity of these mutants when grown on plates (wild-type 91-100%, *spt6-YW* 77-93%, *spt6-50* 72-99% viability). We also note that, for reasons we don’t understand, the integration of the genes necessary for BrdU uptake and incorporation (hENT1 and HSV-1 TK; (Lengronne et al., 2001)) caused a growth defect specifically in *spt6-50* strains, increasing their generation time from 2.5 hours to 3.8 hours (compared to 1.5 hours for wild-type).

**Figure 2.**
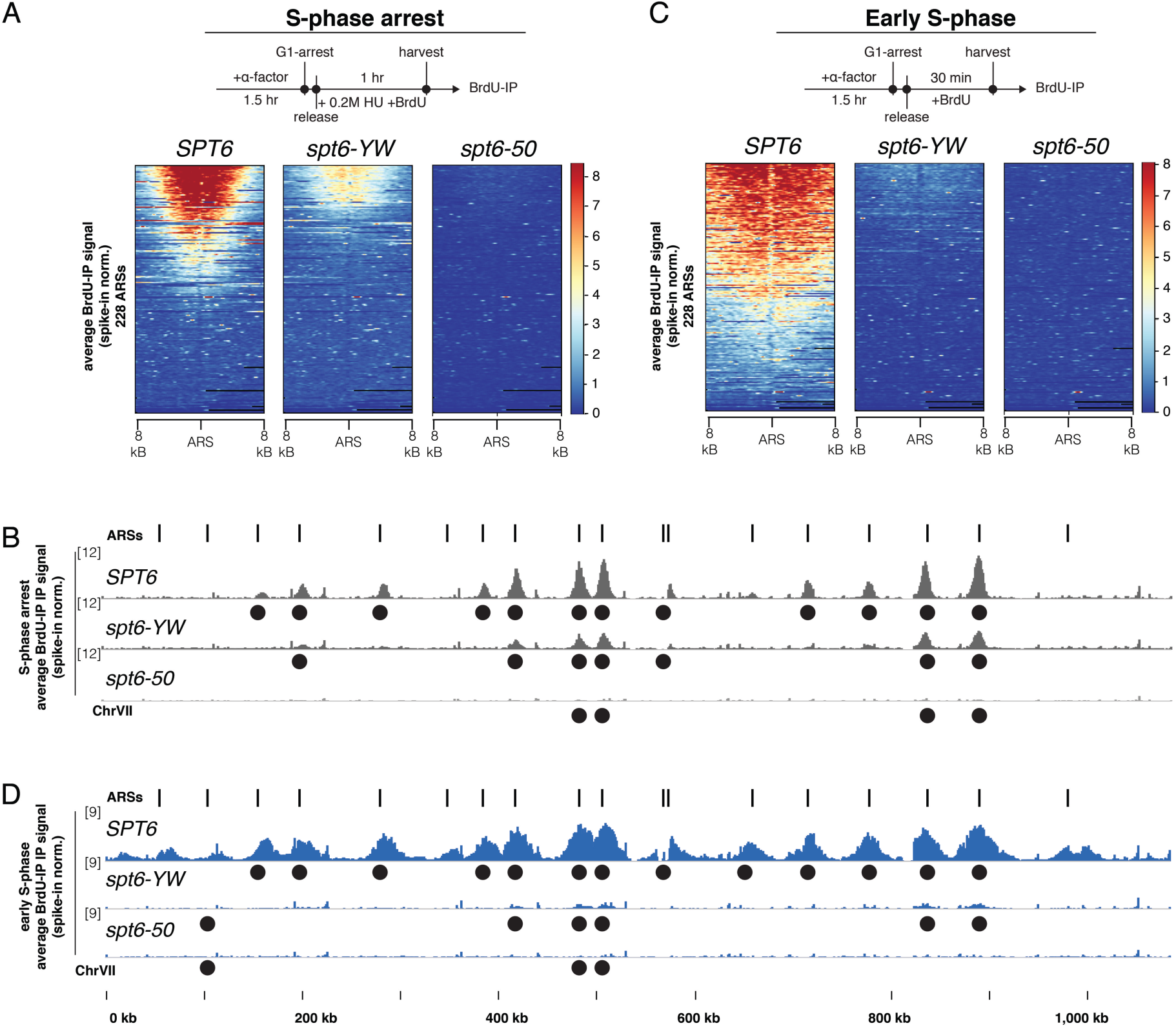
*spt6* mutants have DNA replication defects *in vivo.* (A,C) Spike-in normalized BrdU-IP-seq signal at 228 origins (ARSs) in indicated strains and growth conditions. (B,D) A representative genome browser view of *S. cerevisiae* chromosome VII. Previously identified ARSs are indicated by the vertical lines above. Black circles represent ARSs called in our peak-calling pipeline. The results for S-phase arrested cells (gray, B) and early S-phase cells (blue, D) are shown.

To analyze our results, we first plotted the spike-in normalized BrdU-IP signal for wild-type and *spt6* mutants at a set of defined 228 *S. cerevisiae* replication origins (Feng et al., 2006; Nieduszynski et al., 2006; Raghuraman et al., 2001). For the wild-type strain, BrdU incorporation occurred at a subset of the origins, as expected for early S phase arrested cells (Figure 2A). These were primarily early origins based on previously reported replication timing information (Hawkins et al., 2013) (Figure S2B). For the two *spt6* mutants, we observed a striking decrease in the number of origins that were activated and the level of BrdU incorporation at those origins, with incorporation in the *spt6-50* mutant barely detectable (Figure 2A, B).

To specifically identify which origins were activated across samples, we calculated the overlap between peaks of BrdU signal and the 228 origins (see Methods). In wild-type cells we identified BrdU incorporation at 97 of the 228 origins. In contrast, in *spt6-YW* we detected only 50 origins and in *spt6-50* we detected only 37 origins. An example of origins that are activated compared to the BrdU-IP signal is shown for chromosome VII, one of the largest chromosomes (Figure 2B). All but one detected origin in each *spt6* mutant was in the subset of the 97 origins identified in wild-type cells (Figure S2D). Our results indicate that a subset of origins is activated in each of the *spt6* mutants.

As HU treatment might confound our analysis by inducing replication stress or by reducing the level of RNAPII on chromatin (Poli et al., 2016), we repeated our BrdU-IP without HU treatment. We released alpha-factor synchronized cells into fresh medium and harvested cells after 30 minutes (Figure 2C). Based on flow-cytometry data, this time point represents early S phase cells, similar to the block imposed by HU. Again, duplicate biological replicates were collected and they correlated well, with the Pearson correlations for spike-in normalized IP signal at all loci ranging from 0.86 – 0.98 (Figure S2C).

Our results from these samples were similar to our results from the HU-treated samples in that many fewer origins were activated in *spt6* mutants compared to wild-type (Figure 2C). In the early S phase wild-type samples, we identified 114 origins, a greater number compared to the HU treated samples, showing that the 30-minute time point captures cells later in S phase than does HU-induced arrest. We identified 45 origins in *spt6-YW* and 12 origins in *spt6-50*, which were largely a subset of peaks observed in wild-type samples (Figure 2D, Figure S2D). Similar to our HU results, we again observed a dramatic reduction in signal at origins in both *spt6-YW* and *spt6-50* (Figure 2C, D).

Together, our results show that Spt6 depletion (Figure 1) and *spt6* mutants (Figure 2) cause delays in DNA replication. As we did not observe aberrant activation of late-firing origins in *spt6* mutants, nor did we observe significant signals at non-origin regions (Figure 2, data not shown), we conclude that *spt6* mutants do not regulate DNA replication initiation fidelity. Rather, our data suggests that initiation and/or elongation is impaired at early replication origins in *spt6* mutants.

### Spt6 Physically Interacts with the DNA Replication Machinery

To assess if Spt6 is directly involved in DNA replication, we tested for a physical interaction between Spt6 and the DNA replication machinery. Specifically, we performed co-immunoprecipitation experiments between Spt6 and two components of the MCM complex, a core member of the CMG replicative helicase involved in DNA replication initiation and elongation (Maiorano et al., 2006).

Our results showed physical interactions between Spt6 and both Mcm2 and Mcm4. First, when we immunoprecipitated either Mcm2 or Mcm4, Spt6 co-immunoprecipitated (Figure 3A, B). Second, when we immunoprecipitated Spt6, we found that both Mcm2 and Mcm4 co-immunoprecipitated (Figure 3C). The Spt6-MCM co-immunoprecipitation was largely resistant to treatment with the endonuclease benzonase (Figure 3A-C), suggesting that the interaction is not simply caused by DNA-mediated interactions between two chromatin-associated factors. We did observe that the Mcm2-Spt6 interaction is more susceptible to benzonase treatment and therefore focused our subsequent analyses on the Mcm4-Spt6 interaction. Overall, these results demonstrated that Spt6 interacts with DNA replication components, supporting a direct role for Spt6 in DNA replication.

**Figure 3.**
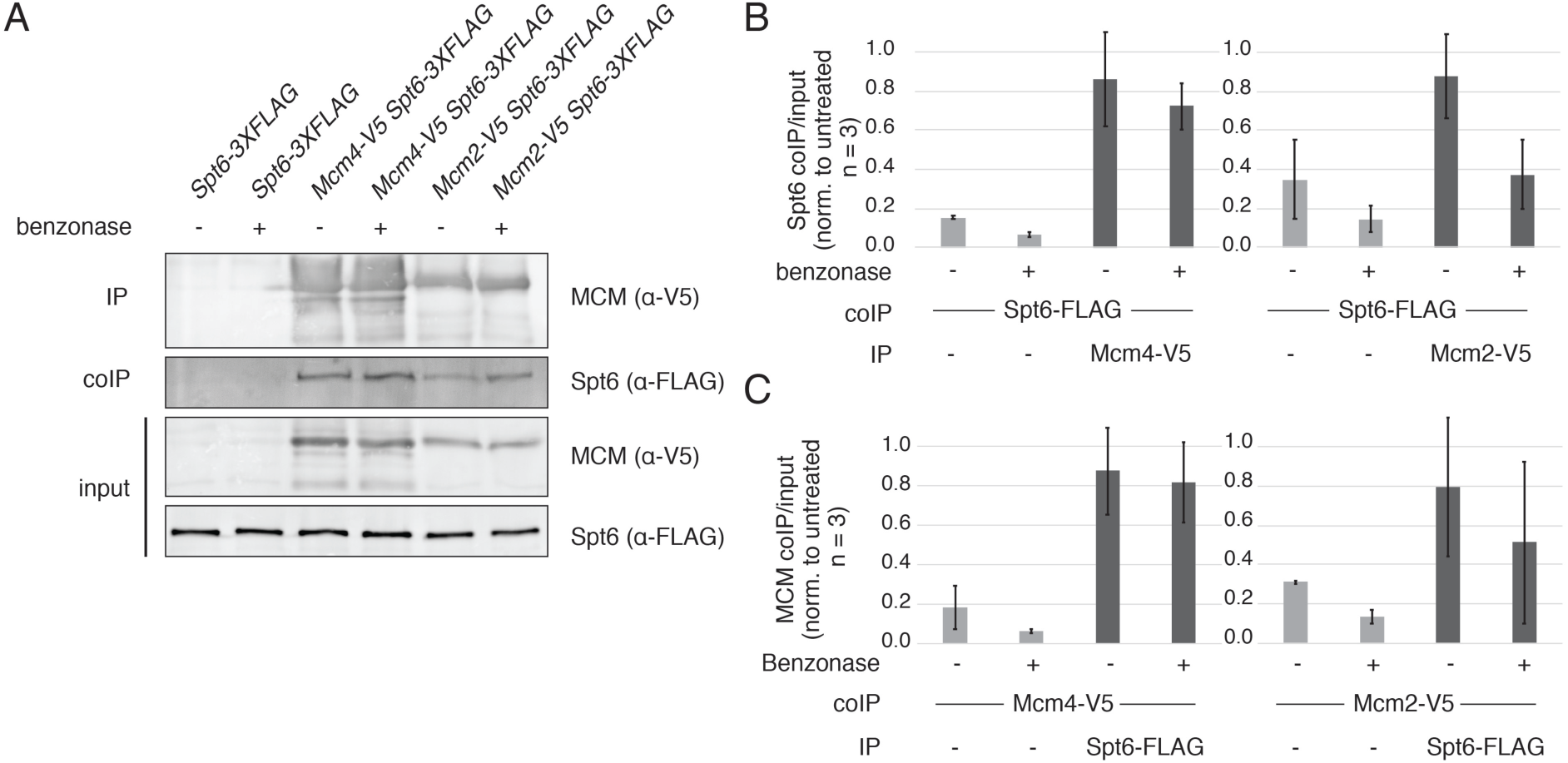
Spt6 physically interacts with the MCM complex. (A) A representative Western blot showing the immunoprecipitation of either Mcm2-V5 or Mcm4-V5 and the co-IP of Spt6-FLAG. Benzonase (100U) was added where indicated. (B) Quantification of three independent coIP experiments as shown in A. (C) Quantification of three independent coIP experiments in which Spt6 was immunoprecipitated and Mcm4 or Mcm2 was assayed for coIP.

### The Spt6 C-terminal SH2 Domains are Required for the Spt6-MCM Interaction

Given the DNA replication defects of the *spt6* mutants, we tested whether the Spt6 mutant proteins had altered interactions with the MCM complex. To test this, we assayed co-immunoprecipitation between the Spt6-YW and Spt6-50 mutant proteins and Mcm4.

Our results show that Spt6-Mcm4 co-immunoprecipitation was greatly decreased in *spt6-50* mutants compared to wild-type, suggesting that the Spt6 SH2 domains are required for the Spt6-MCM interaction (Figure 4A, B). The Spt6-Mcm4 interaction in *spt6-YW* was also mildly impaired; while there was no change in co-immunoprecipitation when Mcm4 was immunoprecipitated, there was a two-fold reduction when Spt6 was immunoprecipitated (Figure 4C).

**Figure 4.**
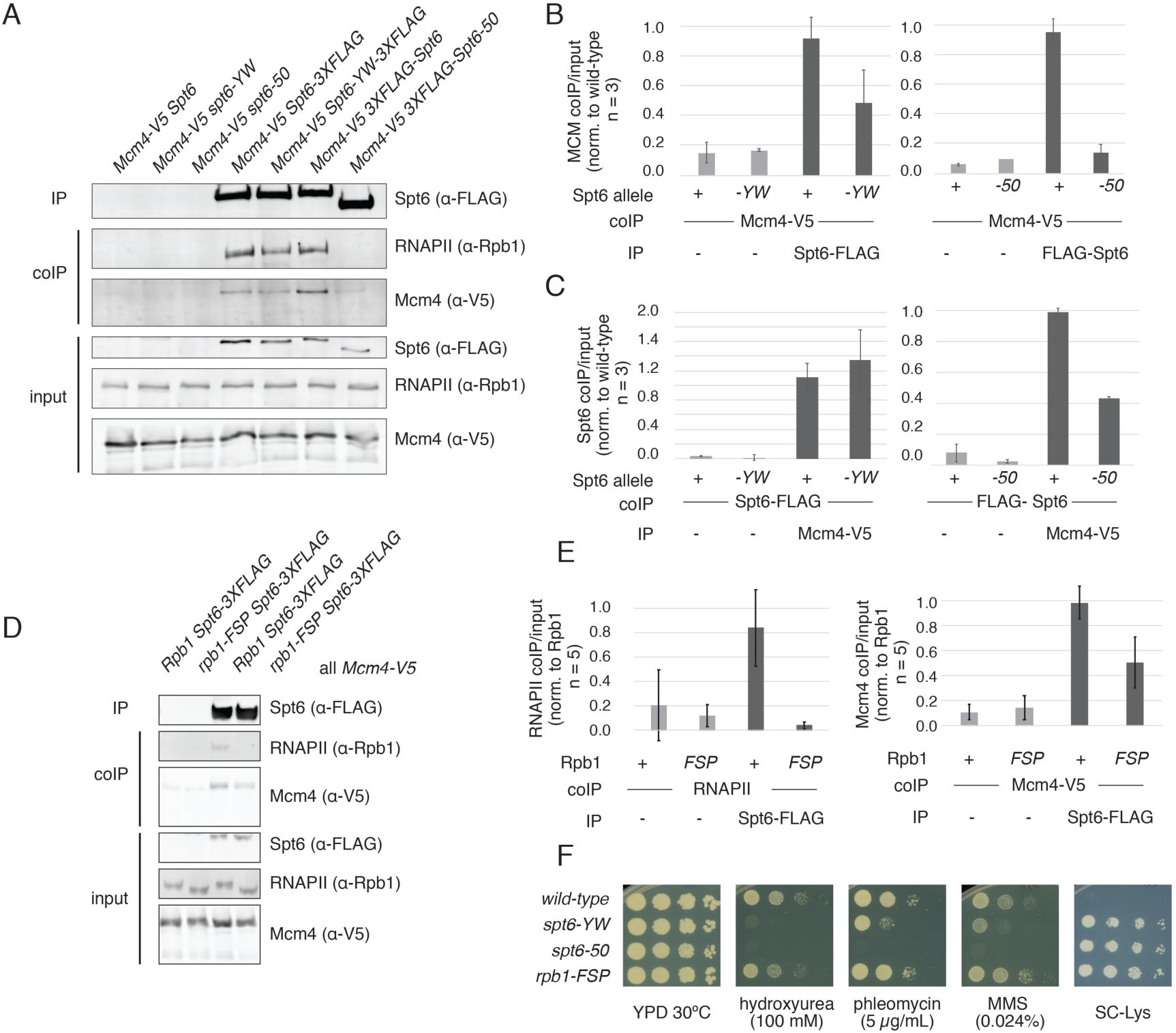
The SH2 domain of Spt6 is required for the Spt6-Mcm4 interaction. (A) A representative Western blot showing the immunoprecipitation of either wild-type or mutant Spt6 and the co-IP of Rpb1 and Mcm4. (B) Quantification of three independent coIP experiments as shown in A. (C) Quantification of three independent coIP experiments in which Mcm4-V5 was immunoprecipitated and wild-type or mutant Spt6 coIP was assayed. (D) A representative Western blot of Spt6 immunoprecipitation in *RPB1* or *rpb1-FSP* strains, in which Mcm4-V5 and Rpb1 coIP where assayed. Quantification of five independent replicates is shown below the blots. (E) Quantification of three independent coIP experiments. (F) Serial dilution plating of *spt6* and *rpb1* mutants. Strains were serially diluted 10-fold, plated on the indicated plates, and incubated at 30°C for three days. All strains contain the *lys2-128δ* reporter.

Several previous studies have shown that the Spt6 SH2 domains are required for the interaction of Spt6 with Rpb1, the largest subunit of RNAPII (Close et al., 2011; Diebold et al., 2010b; Sdano et al., 2017; Sun et al., 2010). Therefore, we tested if the requirement of the SH2 domains for the Spt6-Mcm4 interaction was dependent on the Spt6-RNAPII interaction. We repeated our Spt6-Mcm4 co-immunoprecipitation experiments using a mutant Rbp1*, rpb1-FSP*, that disrupts the RNAPII-Spt6 interaction (Sdano et al., 2017). Our results demonstrate that Spt6-Mcm4 co-immunoprecipitation still occurred, although it was decreased, when the Spt6-RNAPII interaction was disrupted (Figure 4D, E). As expected, the *rpb1-FSP* mutation abolished detectable Spt6-Rpb1 co-immunoprecipitation. These results indicate that at least a portion of Spt6 binds the replication machinery independently from its interaction with the RNAPII machinery. Possibly there is an additional RNAPII-dependent pathway that brings Spt6 to the replication machinery, such as via transcription-replication conflicts.

The *rpb1-FSP* mutation also allowed us to test whether the sensitivity of *spt6* mutants to chemicals that cause replication stress was caused by impairment of the Spt6-RNAPII interaction or of the Spt6-MCM interaction. Our results showed that only the *spt6* mutants, and not the *rpb1-FSP* mutant, were sensitive to compounds that cause DNA replication stress, in agreement with a previous study (Connell et al., 2021). Both classes of mutants exhibited transcription defects, as measured by the ability to grown on SC-Lys plates in the presence of the *lys2-128δ* transcriptional reporter (Figure 4F). Together, these results provide further evidence that Spt6 functions directly in DNA replication.

### *spt6* Mutants Have Negative Genetic Interactions with DNA Replication-Associated Factors

To test for additional genetic evidence for a role for Spt6 in DNA replication, we analyzed *spt6* mutants with respect to altered growth in combination with different classes of mutations that cause replication stress. First, we hypothesized that if Spt6 is involved in DNA replication, *spt6* mutants would exhibit growth defects in combination with replisome mutants. To test this, we focused on Ctf4, an important but nonessential member of the DNA replication machinery. Ctf4 acts as a structural hub within the replisome, linking the replicative helicase, primase, and several other factors to coordinate DNA replication (Gambus et al., 2009; Simon et al., 2014; Tanaka et al., 2009; Villa et al., 2016). We found that, compared to each single mutant, the *spt6-YW ctf4𝛥* and *spt6-50 ctf4𝛥* double mutants had increased growth defects under permissive growth conditions and were extremely sensitive to low doses of DNA replication stress inducing agents (Figure 5A, Figure S3). The severe growth defects of *spt6* mutants in combination with loss of a DNA replication factor provide additional evidence that Spt6 is involved in DNA replication.

**Figure 5.**
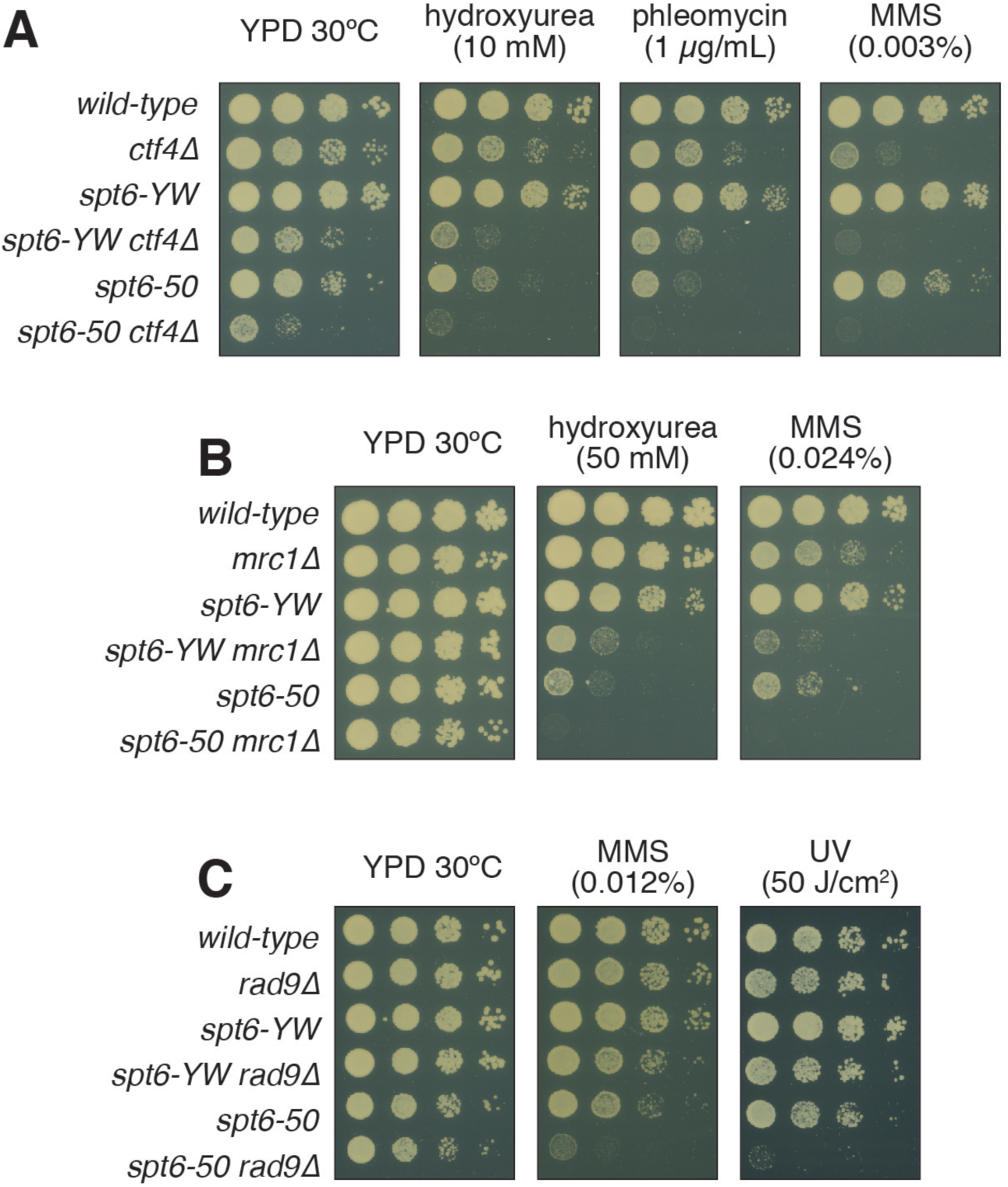
*spt6* mutants have negative genetic interactions with mutations that impair a DNA replication factor and S-phase checkpoint factors. (A-C) Strains were serially diluted 10-fold, plated on the indicated plates, and incubated at 30°C for three days. (A) Ctf4 is a non-essential member of the DNA replication fork complex. (B) Mrc1 is an S-phase checkpoint mediator that responds to stalled DNA replication forks. (C) Rad9 is an S-phase checkpoint mediator that responds to DNA damage.

Second, if *spt6* mutants impair DNA replication, we expect that they would have an increased reliance on S phase checkpoint factors. There are two main branches of the yeast S phase checkpoint. One branch responds to DNA damage, mediated by Rad9, while an alternate branch responds to stalled DNA replication forks, mediated by Mrc1 (Pardo, Crabbé and Pasero, 2017). Our results showed that both *spt6-YW* and *spt6-50* caused increased sensitivity to treatments that induce S phase checkpoint activation when combined with *mrc1*𝛥 (Figure 5B, Figure S3). We observed milder effects when *spt6* mutants were combined with *rad9𝛥,* as only *spt6-50* double mutants had synthetic growth defects (Figure 5C, Figure S3). As a control, we also combined *spt6* mutations with *mad2𝛥*; Mad2 is a spindle-assembly checkpoint factor (Musacchio, 2015). We observed no genetic interactions for these double mutants under the same conditions (Figure S3). The strong negative genetic interactions of *spt6* mutations with *mrc1*𝛥 suggest that stalled DNA replication forks are particularly difficult for *spt6* mutants to overcome in the absence of this S phase checkpoint.

### *spt6* Mutants Have Elevated Levels of DNA Double-Strand Breaks

One consequence of increased replication stress is the formation of DNA double strand breaks (DSBs) (Zeman and Cimprich, 2014). DSBs formed during S phase are primarily repaired by the homologous recombination (HR) pathway (Mathiasen and Lisby, 2014). In yeast, Rad52 is a critical component of the HR repair pathway. As *spt6* mutants had sensitivities associated with replication stress and showed a dependence on S phase checkpoints, we hypothesized that they may also have elevated levels of DSBs.

To measure the level of DSBs in *spt6* mutants, we took advantage of a previous observation that DSBs can be assayed by the presence of Rad52-YFP foci (Lisby et al., 2001). Our results showed that *spt6* mutants have an increased level of Rad52-YFP foci, ranging from a modest effect for *spt6-YW* to a large increase for *spt6-50*. These results show that *spt6* mutants have an elevated level of DSBs (Figure 6A).

**Figure 6.**
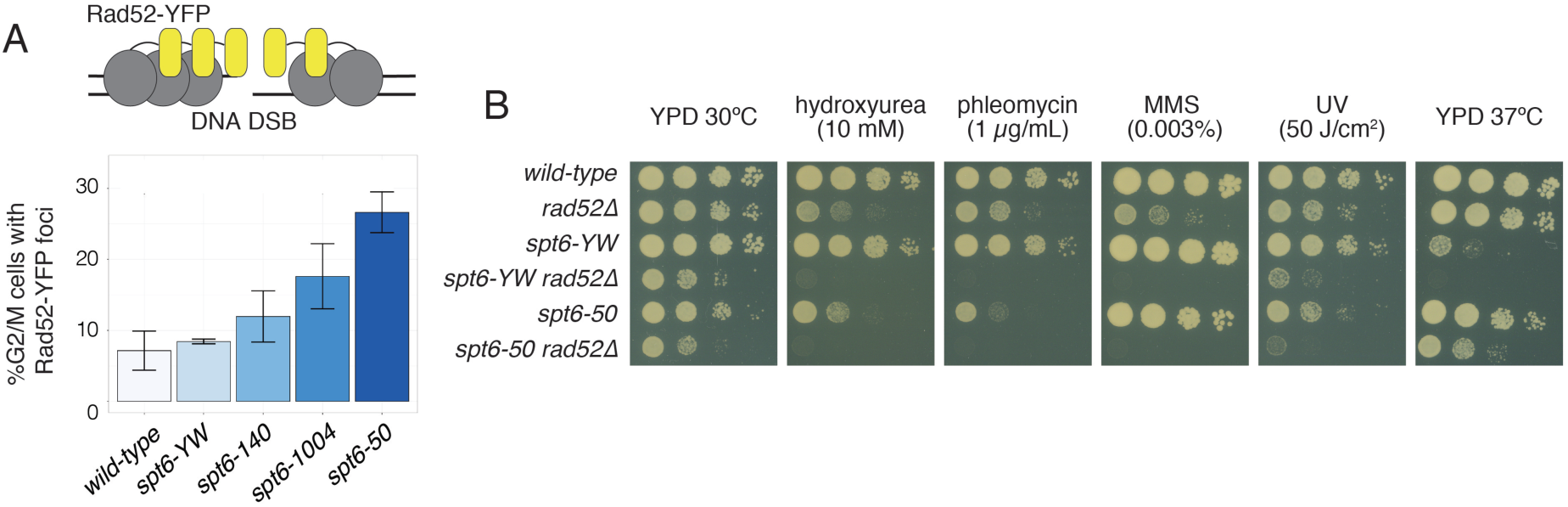
DSB levels are elevated in *spt6* mutants. (A) Quantification of Rad52-YFP foci in G2/M cells. Cells were staged based on cell morphology and the foci were manually counted. Average values are plotted for at least two replicates and over 500 cells per genotype. (B) Analysis of *spt6 rad52Δ* double mutants. Strains were serially diluted 10-fold, plated on indicated plates, and incubated at 30°C for three days.

Given the elevated level of Rad52-YFP foci, we tested for genetic interactions between *spt6* mutations and a deletion of *RAD52*. Our results showed a mild synthetic growth defect for both *spt6-YW rad52*𝛥 and *spt6-50 rad52*𝛥 double mutants compared to the single mutants at permissive and elevated temperatures (Figure 6B). These growth defects were exacerbated when the cells were treated with drugs that induce replication stress. In contrast, we observed no growth effects when *spt6* mutants were combined with a deletion of *lig4*, a member of the non-homologous end joining pathway (Figure S3). This suggests that the increased level of DSBs in *spt6* mutants makes the cells more dependent on homologous recombination.

### Transcription Also Contributes to *spt6*-Mediated Genome Instability

Early studies in yeast suggested Spt6 regulates genome instability (Basrai et al., 1996; Malagon and Aguilera, 1996; Malagón and Aguilera, 2001). Recent work has begun to further characterize these genome instability defects in both yeast and mammalian cells. Through its role in transcription, Spt6 has been shown to cause cell cycle defects (Dronamraju et al., 2018), regulate DSB repair in cancer cells (Obara et al., 2020), and lead to the accumulation of harmful RNA-DNA hybrids at noncoding RNA loci (Nojima et al., 2018). Furthermore, human Spt6 is required for immunoglobulin class-switch recombination in mammalian cells (Begum et al., 2012; Okazaki et al., 2011), a process that requires an induced DNA double strand break, and yeast *spt6* mutants have elevated levels of DSBs (this study). Given that both DNA replication and transcription can independently lead to genome instability (Aguilera and García-Muse, 2013), we addressed whether replication and/or transcription was the underlying cause of Spt6-associated genome instability.

First, we examined hyper-recombination in *spt6* mutants, as it was previously reported that an *spt6-140* mutant is hyper-recombinogenic (Malagon and Aguilera, 1996; Malagón and Aguilera, 2001). Our results showed that all four *spt6* mutants displayed increased levels of recombination compared to wild-type (Figure 7A), from a 3-fold increase for *spt6-1004* to a 30-fold increase for *spt6-50*. Previous studies showed that a mutant in the histone chaperone FACT also caused a 3-fold increase in recombination frequency under similar conditions (Herrera-Moyano et al., 2014). To test if transcription contributes to the hyper-recombination phenotype, we used a second reporter that contains a transcription terminator that prevents transcription across the reporter locus (Figure 7B, Figure S4). Our results showed that inhibiting transcription reduces, but does not abolish the hyper-recombination phenotype for the *spt6-140* and *spt6-50* mutants. This indicates that transcriptional defects contribute in part to Spt6-mediated hyper-recombination in at least some *spt6* mutants, but also suggests that hyper-recombination is also induced by transcription-independent defects, such as in replication.

**Figure 7.**
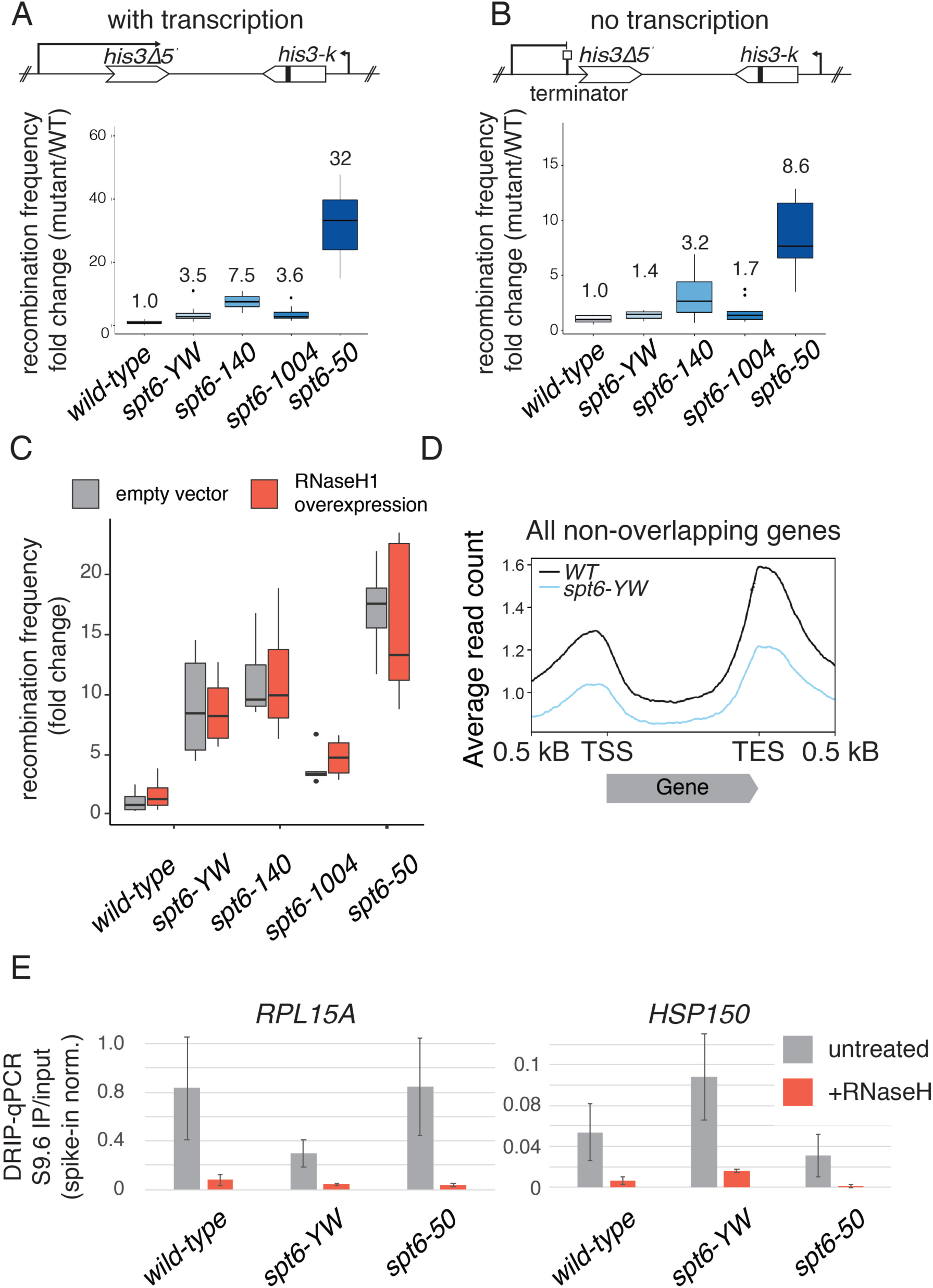
Transcription contributes to *spt6* genome instability. (A) Quantification of an inverted repeat recombination assay (Malagon and Aguilera, 1996). Two independently derived strains and six independent colonies were assayed per genotype. The values represent the average across all samples of a particular genotype. Wild-type values were averaged across replicates. (B) Quantification of inverted repeat recombination assay in the absence of transcription; conducted as in B. (C) Quantification of an inverted repeat recombination assay, as in A except that strains were grown in SC-Trp media and plasmid expression was induced by overnight treatment with estrodiol. (D) Metagene profile of all *S. cerevisiae* non-overlapping genes scaled to their respective transcription start site (TSS) and transcription end site (TES) from DRIP-seq. Data represents the average of two biological replicates. (E) Spike-in normalized DRIP-qPCR data IP/input for two known R-loop containing mRNA genes: *RPL15A* and *HSP150*. The average value for biological duplicates is shown.

### R-loops Are Not a Major Contributor to *spt6*-Mediated Genome Instability

Next, we asked if R-loops are elevated in yeast *spt6* mutants, as a recent study showed that human Spt6 controls R-loop levels over some non-coding regions (Nojima et al., 2018). R-loops are transcription-dependent nucleic acid structures that consist of an RNA-DNA hybrid and an extruded single-stranded DNA loop. They have both regulatory and toxic effects on cells and have been the focus of many recent studies. (Bayona-Feliu and Aguilera, 2021; Castillo-Guzman and Chédin, 2021; Crossley et al., 2019; Kemiha et al., 2021)

If increased RNA-DNA hybrid formation in *spt6* mutants contributed to genome instability phenotypes, we would expect that altering RNA-DNA hybrid levels might enhance or suppress *spt6* mutant phenotypes. To test this, we manipulated the levels of the RNA-DNA hybrid specific endoribonuclease RNaseH in our mutants. First, we combined each of the four *spt6* mutations with deletion mutations for the two catalytic RNaseH enzymes in the yeast genome, *RNH1* and *RNH201.* In these mutants, RNA-DNA hybrid levels are greatly increased (Wahba et al., 2016). Our results showed no change in the *spt6* phenotypes, suggesting that increasing RNA-DNA hybrid levels does not exacerbate the *spt6* genome instability phenotypes (Figure S3). Next, we tested whether reducing RNA-DNA hybrid levels by overexpression of *RNH1* would suppress hyper-recombination. We found that overexpression of *RNH1* did not reduce the recombination frequency compared to expression of an empty vector control in *spt6-YW*, *spt6-140*, or *spt6-1004* strains (Figure 7C). The *spt6-50* mutant had more variable results, although there was no statistical difference compared to wild-type.

Finally, we measured the level of RNA-DNA hybrids in two of our mutants using DNA-RNA hybrid IP (DRIP). In this method, RNA-DNA hybrids are directly immunoprecipitated using the S9.6 antisera. Genome-wide experiments in *spt6-YW* indicated that the level of RNA-DNA hybrids is decreased compared to wild-type. We note that there was no spike-in to normalize for the genome-wide data (Figure 7D). Therefore, we repeated these experiments adding exogenous spike-in *S. pombe* genomic DNA and assayed by DRIP-qPCR both *spt6-YW and spt6-50 (*Figure 7E). We observed variable RNA-DNA hybrid levels across two tested loci in wild-type samples compared to *spt6* mutants. However, we did not observe increased levels of RNA-DNA hybrids in *spt6* mutants. As RNA-DNA hybrid levels correlate with transcription, these findings are in agreement with *spt6-YW* genome-wide RNAPII occupancy data (Viktorovskaya et al., 2021), in which RNAPII levels are also reduced. In conclusion, our results suggest that altered RNA-DNA hybrid levels do not cause the *spt6* genome instability phenotypes.

## DISCUSSION

The results presented in this paper provide strong evidence that Spt6, a histone chaperone previously shown to control transcription and chromatin structure, also functions directly in DNA replication. A functional requirement for Spt6 in DNA replication is shown by our analysis of both Spt6 depletion and *spt6* mutants, which revealed a delayed S phase and reduced DNA synthesis from early origins of replication. Our evidence for a direct role for Spt6 comes from the demonstration of co-immunoprecipitation between Spt6 and the MCM complex, a core member of the eukaryotic replisome. Supporting data also includes extensive genetic analyses that showed the sensitivity of *spt6* mutants to DNA-damaging agents, as well as double mutant analyses that indicated greatly enhanced mutant phenotypes when *spt6* mutations were combined with mutations that impair DNA replication factors, S-phase checkpoint regulators, and recombination factors. Taken together, these results suggest that Spt6, like some other histone chaperones, including Asf1 (Mousson et al., 2007) and FACT (Formosa and Winston, 2021), are required for both DNA replication and transcription.

Our finding that the Spt6 carboxy-terminal SH2 domains are required for the Spt6-MCM interaction raises several interesting questions. The Spt6 SH2 domains were previously shown to bind to Rpb1 of RNAPII, dependent upon Rpb1 phosphorylation at specific sites (Close et al., 2011; Diebold et al., 2010a; Sdano et al., 2017; Sun et al., 2010; Yoh et al., 2007). Therefore, it is conceivable that the Spt6-replisome interaction also requires specific phosphorylation of MCM or other replisome proteins. Phosphorylation of the replication machinery is a critical regulatory step that activates “licensed” replication origins upon entry into S phase and initiates DNA replication (Labib, 2010). Such temporal specificity would suggest that Spt6 is required for a particular step in DNA replication. In addition, our results demonstrated that when the Spt6-RNAPII interaction is disrupted by mutation of RNAPII, the Spt6-MCM interaction still occurs, although it is reduced. This reduction suggests a role for the Spt6-RNAPII interaction in facilitating an Spt6-MCM interaction.

Possibly, Spt6 is required to resolve transcription-replication conflicts, either independently or as part of a histone chaperone network. Interestingly, a recent study showed that Spt2, another histone chaperone, is required to help resolve transcription-replication conflicts (Zardoni et al., 2021). As the recruitment of Spt2 to chromatin is partially dependent upon Spt6 (Nourani et al., 2006), this suggests one mechanism by which Spt6 might maintain genome stability. Another intriguing aspect of our results relates to the close functional connections between Spt6 and FACT (van Bakel et al., 2013; Cheung et al., 2008; Jeronimo et al., 2015; McCullough et al., 2015; Viktorovskaya et al., 2021), an essential histone chaperone that functions in transcription, replication, and mediates transcription-replication conflicts (Duina, 2011; Formosa and Winston, 2021; Herrera-Moyano et al., 2014). The respective contributions of Spt6 and FACT to DNA replication and genome stability are interesting topics for future studies, especially as FACT emerges as a potential therapeutic target in cancer (Gasparian et al., 2011).

Our results for Spt6 in yeast, along with a recent result from studies of Spt6 in human cells (Narain et al., 2021), have shown that Spt6 is required for DNA synthesis *in vivo*. Based on the previously reported functions of Spt6 in transcription, we hypothesized that Spt6 may contribute to DNA synthesis by stimulating initiation or elongation of DNA polymerases, analogous to how Spt6 facilitates transcription by RNAPII (Ardehali et al., 2009; Endoh et al., 2004; Narain et al., 2021; Nojima et al., 2018; Žumer et al., 2021), or by regulating the fidelity of origin activation, similarly to how Spt6 regulates the transcription of thousands of cryptic intragenic transcription start sites (Cheung et al., 2008; Doris et al., 2018; Gouot et al., 2018; Kaplan et al., 2003; Uwimana et al., 2017). Our BrdU-seq analysis showed that the defect in *spt6* mutants occurs by reduced synthesis from known origins, a result consistent with defects in either initiation or elongation, rather than ectopic synthesis from non-origin loci. This difference may reflect the different sequence requirements for the two events, with less specific information required for the initiation of transcription versus the initiation of DNA replication, which is defined by the ARS sequence in *S. cerevisiae* (Nieduszynski et al., 2006). It would be interesting to see if loss of Spt6 function resulted in cryptic replication in mammalian cells, where DNA origins are not defined by known DNA binding motifs (Lee et al., 2021).

Histone chaperones contribute to genome instability through a variety of different mechanisms. One mechanism that has been the focus of recent studies, is the relationship between histone chaperones, such as FACT (Cristini et al., 2018; Herrera-Moyano et al., 2014), and R-loops. A recent study depleted Spt6 from mammalian cells and demonstrated that the abundant R-loop signatures at mRNA loci were decreased genome-wide, while the small proportion of R-loops present at non-coding loci increased (Nojima et al., 2018). These changes correlated with the level of transcription present at each transcription unit, as has been previously seen (Sanz et al., 2016). Our finding, that yeast *spt6* mutants do not have elevated levels of R-loops, agrees with this work. It further highlights that the replication stress observed in *spt6* mutants, or depletion of Spt6 from mammalian cells, may be caused by a direct role of Spt6 in replication or a role for Spt6 in mediating transcription-replication conflicts.

In conclusion, our studies have unveiled a role for Spt6 in DNA replication. Further work is needed to define the precise interactions of Spt6 with the replisome and to continue to understand the connection between the roles of Spt6 in replication, transcription, and conflicts between the two. Future studies will benefit from the identification of *spt6* alleles that specifically impair replication and not transcription, as such alleles have been valuable in studies of both FACT (Yang et al., 2016) and another factor that controls both transcription and replication, Sen1 (Appanah et al., 2020).

### Limitations of the study

Our study reveals an import function of Spt6 in DNA replication. However, to fully demonstrate a role of Spt6 in DNA replication, *in vitro* DNA replication assays with purified components should be performed. Another limitation of our study is the growth defect that occurs in the *spt6-50* mutant when combined with BrdU incorporation components. This could lead us to over-estimate the effect of the *spt6-50* mutant on DNA replication as measured by BrdU-IP. We note that the *spt6-50* mutant used in co-immunoprecipiation experiments does not have this defect. Finally, while we have tried to separate the transcription and replication defects observed in *spt6* mutants, true separation of function alleles are still lacking and could further differentiate direct and indirect mutant phenotypes.

## ACKNOWLEDGMENTS

We thank Kaia Mattioli and James Warner for helpful comments on the manuscript. We also thank Lorenzo Costantino and Doug Koshland for generously supplying S9.6 antisera, Etienne Schwob for supplying plasmids and advice for the BrdU experiments, and Nick Rhind for supplying *S. pombe* strains for BrdU experiments. We are also grateful to Marco Fumasoni for guidance on the flow cytometry experiments and Jessica King for help with experiments. Part of this research was conducted on the O2 High Performance Computer Cluster supported by the Research Computing Group at Harvard Medical School. This work was supported by grants from the NSF and from the Landry Cancer Biology Consortium to C.L.W.M. and NIH Grant R01GM135251 to F.W.

## AUTHOR CONTRIBUTIONS

C.L.W.M. and F.W. designed the experiments. C.L.W.M. performed all of the experiments.

C.L.W.M. and F.W. wrote the manuscript.

## STAR METHODS

### RESOURCE AVAILABILITY

#### Lead contact

Further information and requests for resources and reagents should be directed to and will be fulfilled by the lead contact, Fred Winston.

#### Materials availability

Yeast strains and plasmids generated in this study are available upon request.

#### Data and code availability

- Genome-wide sequencing data have been deposited at GEO and are publicly available as of the date of publication. Accession numbers are listed in the key resources table. All data reported in this paper will be shared by the lead contact upon request.
- All original code has been deposited at GitHub and is publicly available as of the date of publication.
- Any additional information required to reanalyze the data reported in this paper is available from the lead contact upon request.

### EXPERIMENTAL MODEL AND SUBJECT DETAILS

#### Yeast strains and growth

All *S. cerevisiae* strains used in this study are in the S288C background. They were constructed by standard methods either by transformation, cross, or CRISPR-Cas9 mutagenesis (Laughery et al., 2015). All *S. cerevisiae* liquid cultures were grown in YPD (1% yeast extract, 2% peptone, 2% glucose) at 30°C unless otherwise mentioned. For spike-in normalization, *S. pombe* liquid cultures were grown in YES (0.5% yeast extract, 3% glucose, 225 mg/l each of adenine, histidine, leucine, uracil, and lysine) at 32°C.

### METHOD DETAILS

#### Yeast dilution plating (spot tests)

Yeast strains were serially diluted 10-fold and plated onto the indicated media: YPD 30°C (permissive), YPD 37°C (high temperature), hydroxyurea (HU), phleomycin, methyl methanesulfonate (MMS), ultra-violet light (UV), or SC-Lys. Plates were incubated for three days. All experiments were performed at least in duplicate. Each plate contained a wild-type and single mutant control, although only one representative example of these controls is shown.

#### Recombination assays

Recombination frequency reporters were previously described (Malagón and Aguilera, 2001). Recombination frequencies were determined as the average frequency of six independent colonies. For each strain, two independent experiments were performed, for a total of 12 independent colonies per strain. For each colony, strains were plated on SC complete to determine cell viability and SC-His to determine His+ recombinants. Recombination frequencies are reported as the ratio of these two values. Yeast *RNH1* was cloned into pRS414 under the expression of an estradiol-inducible promoter (pKW26). Strains were transformed with pRS414 or pKW26, and grown under selective conditions SC-Trp. RNaseH overexpression was induced by addition of estrodiol overnight. Recombination frequencies were determined as described above, except only six colonies were assayed per genotype.

### DNA-RNA IP (DRIP)

Yeast strains were harvested during mid-log phase (OD∼0.6). For DRIP-seq, genomic DNA was isolated using Qiagen Tip kit according to manufacturer’s protocol with the exception that RNase was not added to buffer G2. Genomic DNA was first treated with S1 nuclease for 30 minutes at 50°C (as described in (Wahba et al., 2016)). Samples were then fragmented by sonication to ∼300 bp. For qPCR analysis, genomic DNA was isolated by phenol:chromoform extract followed by a brief treatment with RNaseA/T1 in 300 mM NaCl. DNA was fragmented by overnight digestion at 37°C with restriction enzymes (AccI, EcoRV, NcoI, HaeIII, BsrG1) in cutsmart buffer in the presence or absence of 20 U RNaseH. For DRIP-seq and DRIP-qPCR, following fragmentation, samples were purified by phenol:chloroform extraction then ethanol precipitated. Samples were resuspended in 1XFA buffer (1% TritonX-100, 0.1% sodium deoxycholate, 0.1% SDS, 50 mM HEPES, 150 mM NaCl, 1 mM EDTA). Antibody S9.6 (25µg) was pre-bound to 100 µL Protein A Dynabeads for 1 hour at 4°C in 1XFA buffer. 100 µg fragmented DNA was incubated with the S9.6-protein A beads for 2 hours at 4°C. Beads were washed in 5 successive washes: 1XFA, 1XFA with 0.5M NaCl, LiCl wash buffer (0.25M LiCl, 0.5% NP-40, 0.5% sodium deoxycholate, 1 mM EDTA, 10 mM Tris-HCl), twice with TE buffer. DNA was eluted in 300 µL elution buffer (1%SDS, 0.1M NaHCO3) for 2 hr at 65°C. Samples were then treated with 1 µL Proteinase K (20 mg/mL) for 1 hour at 42°C, extracted with phenol:chloroform, and then ethanol precipitated. The final DNA pellet was resuspended in 50 µL TE buffer. The percent hybrid signal for input and IP samples were quantified by qPCR. Sequencing libraries were prepared using the Qiagen GeneRead kit according to manufacturer’s instructions. For spike-in normalized qPCR samples, *S. pombe* genomic DNA was prepared as described above and was added after RNase treatment in equal amounts.

#### Alpha-factor synchronization

Yeast strains were grown to an OD∼0.2 and then synchronized by treatment with alpha factor (5 nM for *bar1𝛥* or 5 µM for *BAR1*) for up to 2 hours. Visual inspection confirmed >90% of cells had shmoo-like morphology. Cells were then washed three times with YPD media lacking glucose supplemented with fresh 50 µg/mL pronase. Cells were released into indicated media and harvested as described below.

#### Flow cytometry cell cycle analysis

Following alpha-factor arrest (see above), cells were released into fresh YPD supplemented with 50 µg/mL pronase. Yeast samples were harvested at the indicated time points and fixed in 1 mL 70% EtOH overnight at 4°C. Fixed cells were washed once in 50 mM Tris-HCl pH7.5, then treated with RNaseA overnight at 37°C. Following removal of RNaseA, cells were treated with Proteinase K and incubated at 37°C for 2 hours. Cells were resuspended in 50 mM Tris-HCl pH7.5. Cells were briefly sonicated and then treated with 1 mL 50 mM Tris-HCl pH7.5 with 1µL of 1mM SYTOX green in DMSO. Flow cytometry analysis was immediately performed (Fortessa, BD Biosceince). Data was analyzed with FlowCytometryTools (https://eyurtsev.github.io/FlowCytometryTools/).

#### BrdU-IP

Following alpha-factor arrest (see above), cells were released into fresh YPD supplemented with 50 µg/mL pronase and 400 µg/mL BrdU. For HU-treated samples, hydroxyurea was added to a final concentration of 0.2M HU. Cells were harvested after 30 minutes (untreated) or 60 minutes (HU treatment) by the addition of sodium azide to a final concentration of 0.1%. Cell pellets were washed once with cold 1XTBS and stored at -80°C. Cell pellets were resuspended in lysis buffer (2% SDS, 2% Triton-X100, 100 mM NaCl, 10 mM Tris pH 8.0, 1 mM EDTA) and 500 µL acid washed glass beads were added. Cells were lysed by vortexing for a total of 30 minutes at 4°C. Genomic DNA was isolated by phenol-chloroform isolation followed by ethanol precipitation with NaCl salt. DNA pellet was washed once with 70% ethanol and then resuspended in sterile water. DNA was treated with RNaseA/T1 for 30 minutes at 37°C followed by Proteinase K for 30 minutes at 50°C. Genomic DNA (gDNA) was column purified by a PCR purification kit. gDNA was fragmented by sonication to ∼300 bp. Sonication efficiency was confirmed by gel electrophoresis. DNA concentration was measured by nanodrop and an equal amount of sonicated *S. pombe* genomic DNA was added to each sample. Input samples were removed immediately following spike-in addition. For next-generation sequencing experiments, libraries were prepared with the Qiagen GeneRead kit according to manufacturer’s instructions prior to IP. For the IP, gDNA was boiled for 5 minutes at 95°C. Samples were incubated on ice for 3 minutes. 50 µL of gDNA (50-500 ng in EB) was combined with 50 µL anti-BrdU (1:250 in 2X IP buffer (2XPBS, 0.001% Triton-X100)) and incubated on a rotisserie for 2 hours at 4°C. 15 µL of washed Protein G Dynabeads was added. Samples were incubated on a rotisserie for 1 hour at 4°C. Samples were washed thrice in 1XIP buffer. Sample was eluted by the addition of TES and incubating for 10 min at 65°C. Samples were column purified prior to qPCR or sequence analysis.

#### Co-immunoprecipitation

For co-immunoprecipitation experiments, we used two epitope-tagging schemes. For Spt6 (Figure 3, 4) and Spt6-YW (Figure 4), we a C-terminal 3XFLAG epitope tag. For *spt6-50,* a nonsense mutation resulting in a C-terminal truncation of the protein, we used N-terminally tagged 3X-FLAG-Spt6 and 3X-FLAG-Spt6-50 (Figure 4). Yeast strains were harvested during mid-log phase (150 mL, OD∼0.6). Cell pellets were washed once and stored at -80°C. Cell extracts were prepared by resuspending cell pellets in IP buffer (10% glycerol, 100 mM HEPES, 2mM EDTA, 150 mM KOAc, 0.1% NP-40, 10 mM MgOAc, 2 mM NaF, supplemented with 1 mM DTT and SIGMAFAST protease inhibitor tablet (Sigma) at time of use; (Gambus et al., 2006)) and bead-beating for a total of 6 min at 4°C, with 5 minutes incubation on ice after every minute. For Benzonase treatment, 300U of Benzonase was added for 30 minutes rotating at 4°C. For all samples, cell lysate was then cleared for 20 minutes, 12500 rpm, 4°C. Total protein concentration was measured by Pierce BCA Protein Assay Kit (Thermo Fisher). For V5 immunoprecipitation, 5 µg anti-V5 (Invitrogen) was pre-bound to 30 µL Protein G Dynabeads for 1 hour at 4°C in 1XIP buffer. For FLAG immunoprecipitations, anti-FLAG-M2-FLAG affinity gel (Sigma A2220) was washed and resuspended in 1XIP buffer. An equal concentration of total protein (10-20 µg/µL) was incubated with antibody-beads for 2 hours, rotating, 4°C. Samples were washed thrice in IP buffer and were eluted in 30 µL 3X protein loading dye (9% SDS, 10% BME, 187.5 mM Tris-HCl, 30% glycerol, bromophenol blue) at 105°C for 3 minutes. The beads were then spun down and the supernatant was transferred to a new tube and either stored or used for western blots (6% SDS-PAGE gels). The following antibodies were used for western blots analysis: 1:5,000 anti-FLAG-M2 (Sigma), 1:5,000 anti-V5 (Invitrogen), 1:2,500 anti-RNAPII-CTD (8WG16). Licor secondary antibodies were used. Images were quantified with ImageStudioLite. Quantification was normalized to the signal for the wild-type tagged sample for each blot.

#### Rad52-foci microscopy

Cells were grown to mid-log phase. Samples were washed once with 1XPBS, briefly sonicated, and placed on a 2% agarose pad on imaging slide. Images were acquired with a Nikon Elipse Ti2 using 60X lens. Cells were manually staged based on cell morphology and foci were manually counted.

#### Bioinformatics analysis

A custom pipeline was used to process both BrdU-IP-seq and DRIP-seq data. The custom pipeline was generated using the workflow manager Snakemake (Köster and Rahmann, 2012). Briefly, data was aligned to a combined *S. cerevisiae + S. pombe* genome (*S. cerevisiae* R64-2-1, *S. pombe* ASM294v2) using Bowtie2 (Langmead and Salzberg, 2012). Sequences were trimmed with cutadapt and processed using samtools (Li et al., 2009), bedtools, and deeptools (Ramírez et al., 2016) to generate genome coverage tracks. This pipeline is available on the Winston lab GitHub (https://github.com/winston-lab). Deeptools was used to generate heatmaps of signal at desired regions and MACS2 (Zhang et al., 2008) was used to call peaks on individual samples. For HU-treated BrdU-IP samples, irreproducible discovery rate (IDR) analysis was performed to identify common peaks across samples. As peaks were much broader and weaker for 30-minute samples, IDR analysis was not feasible. Instead, for 30-minute BrdU-IP samples, MACS2 peaks per sample were first filtered to isolate peaks larger than 1kB in length and then merged to combine peaks within 5 kB. Peak regions present in both samples were then used as reproducible peaks across samples.

#### Northern Blotting

RNA extraction from *S. cerevisiae* was done using hot acid phenol extraction. Ten µg of RNA was resuspended in RNA loading buffer (6% formaldehyde, 1X MOPS, 2.5% Ficoll, 10 mM Tris-HCl pH 7.5, 10 mM EDTA, 7 µg/ml ethidium bromide, 0.025% bromophenol blue, 0.025% Orange G), heated at 65°C for 10 minutes, then loaded on a 1% agarose/formaldehyde gel. The gel was run for 400 volt hours. Samples were denatured in the gel in 0.05M NaOH for 30 minutes at room temperature followed by neutralization in Tris-HCl, pH 7.5 for 30 minutes at room temperature. Samples were transferred to Genescreen membrane by upward capillary transfer in 1XSCC solution. The membrane was UV-crosslinked prior to hybridization. The membrane was incubated in pre-hybridization solution (50% deionized formamide, 10% dextran sulphate, 1M NaCl, 0.05M Tris-HCl pH 7.5, 0.1% SDS, 0.1% sodium pyrophosphate, 10X Denhardts reagent, 500 µg/mL denatured salmon sperm DNA) at 42°C for 5 hours. Membrane was hybridized to 32P-dATP labeled probe overnight at 42°C. The membrane was washed: 2 washes with 2XSCC for 15 minutes at room temperature, 2 washes with 2XSCC+0.5% SDS for 30 minutes at 65°C, and 2 washes with 0.1X SCC for 30 minutes at room temperature. Blots were imaged with a Typhoon phosphor-imager.

### KEY RESOURCES TABLE

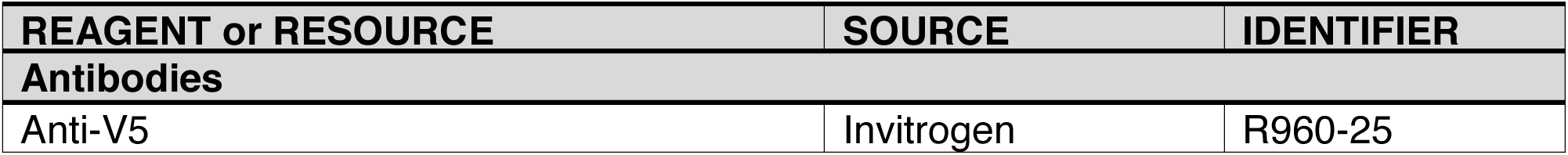
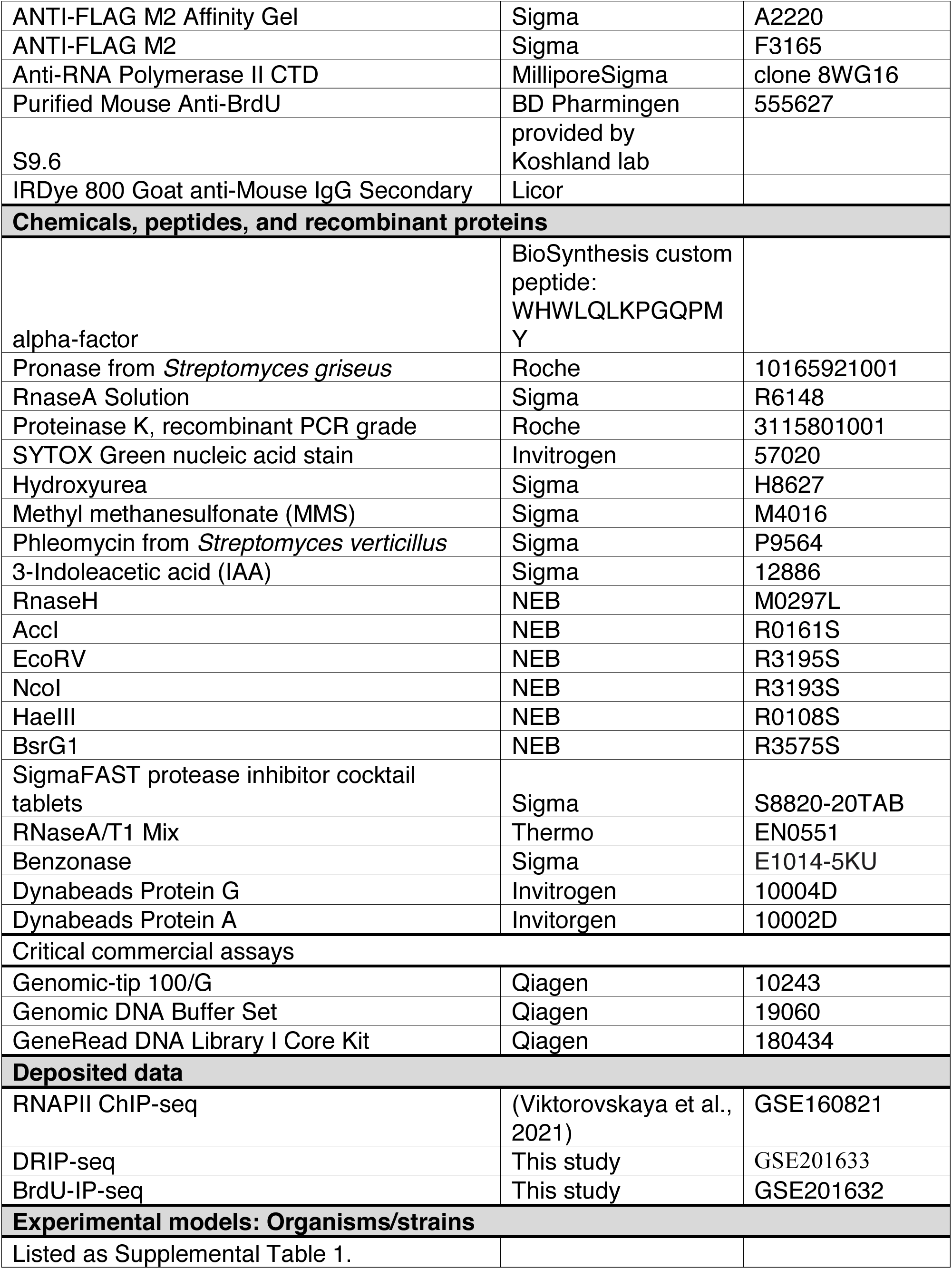
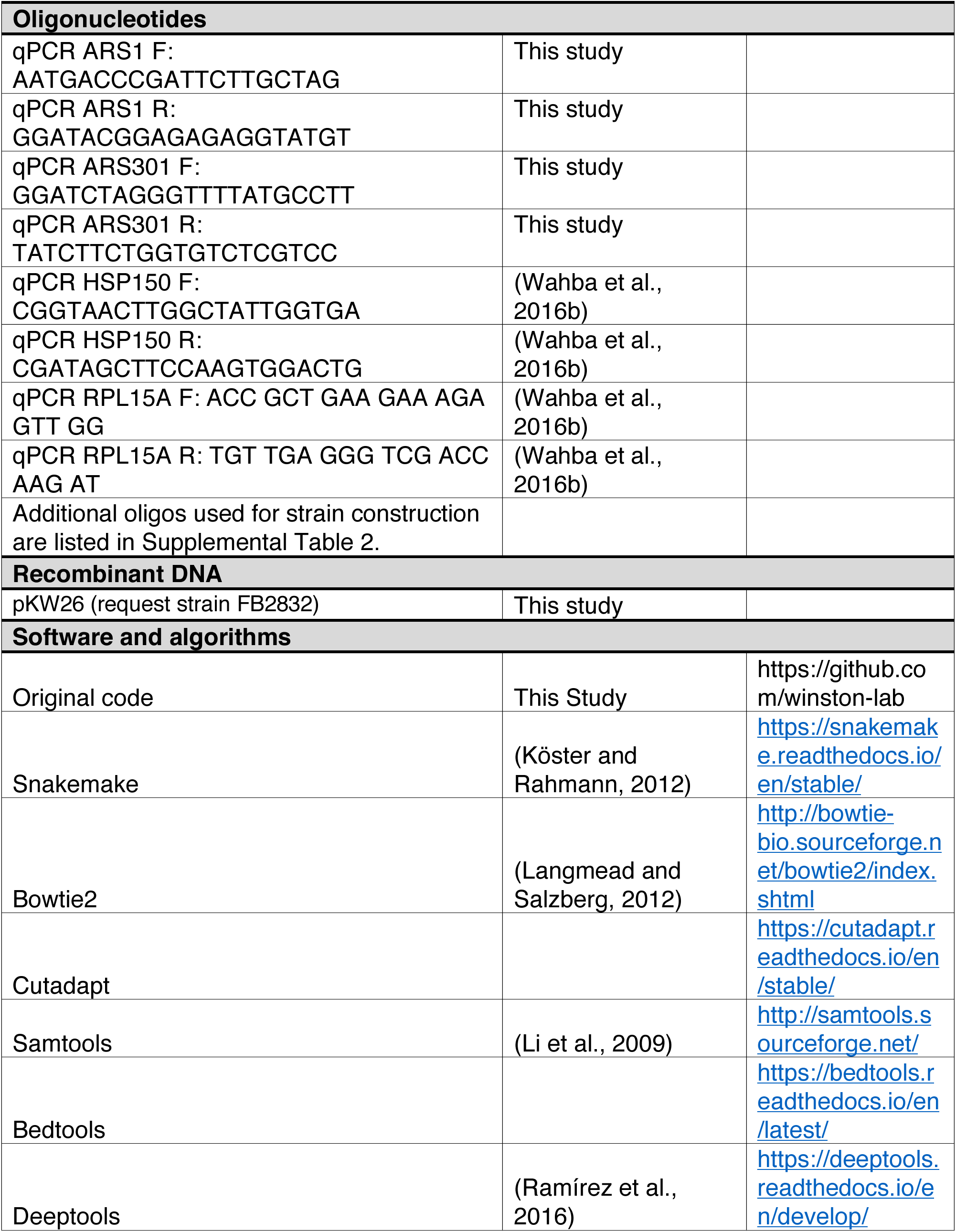
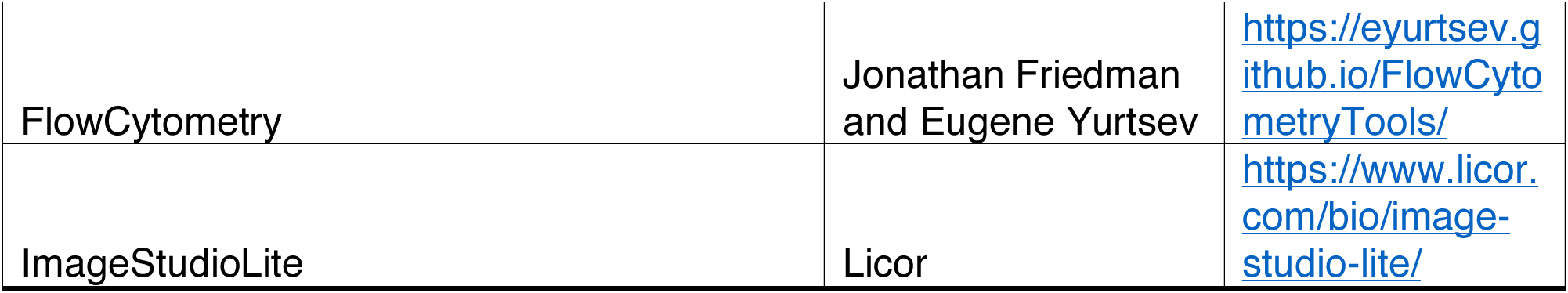

## SUPPLEMENTAL LEGENDS

**Supplemental Table 1.** Related to STAR Methods. Yeast strains used in this study.

**Supplemental Table 2.** Related to STAR Methods. Oligonucleotides used for yeast strain construction.

**Supplemental Figure 1** Related to Figure 1 (A) A representative Western blot of Spt6-AID depletion. Spt6 levels are shown in asynchronously dividing samples (async.), after alpha-factor arrest in the presence of DMSO (D) or IAA (I) for one hour, and after release into fresh media for 90 minutes in the presence of DMSO or IAA. Pgk1 was used as a loading control. Quantification represents the average of biological triplicates. (B) Representative smoothed flow cytometry histogram for indicated time points and conditions. (C) A representative Western blot of the Spt6 protein levels for each *spt6* mutant studied. Quantification of Spt6 protein levels in *spt6* mutants. The average value of three biological replicates is shown. (D) TSS-seq data of *spt6-YW* vs. wild-type for strains grown at 30°C (Viktorovskaya et al., 2021). The values for each gene are plotted based on their mean expression vs the log2(fold change) of *spt6-YW* / wild-type. Differentially expressed genes, defined as having a log2FC > 2, are indicated in black. Genes in red represent SBF targets (Iyer et al., 2001). (E) Shown are the log2 fold change (log2FC) and log adjusted p-values (logpadj) for select genes of interest – including cell cycle regulators, *RNR* genes, and the only SBF targets that are differentially expressed (*GFA1*, *PMA1*).

**Supplemental Figure 2** Related to Figure 2 (A) Pearson correlation plots for individual BrdU-IP spike-in normalized IP signals for S-phase arrest (HU treatment) samples. (B) Replication origins were divided into quartiles based on previously reported replication timing (Hawkins et al., 2013), with an equal number of origins per group. The average value across all 58 ARSs per group is plotted for BrdU-IP S-phase arrested (HU treatment) samples (red for wild-type, blue for *spt6-YW*, and green for *spt6-50*). (C) Pearson correlation plots for individual BrdU-IP spike-in normalized IP signals for early S-phase (30 minute) samples. (D) Comparison of identified activated ARSs across genotypes. Genotypes are indicated on the left with the total number of activated ARSs in parenthesis. Samples are then compared in sets with the number of ARSs activated in that set indicated below the diagram.

**Supplemental Figure 3.** Related to Figure 5. Summary of *spt6* genetic interactions. Indicated strains were spotted on indicated plates. Growth is indicated, ranging from ++++ indicating growth comparable to a wild-type strain to – indicating no detectable growth. Plates were scored on Day 2 or Day 3.

**Supplemental Figure 4.** Related to Figure 7. (A) Average values for all experiments in Figure 7A and 7B. FC indicates fold change. (B) Schematic of reporters. (C) Representative Northern blot analysis of hyper-recombination reporter (Malagón and Aguilera, 2001).

