## Supplemental files for "The conserved histone chaperone Spt6 facilitates DNA replication and mediates genome instability"

Supplemental Table 1. Related to STAR Methods. Yeast strains used in this study.

| Strain Name | Mate Type | Genotype | Reference | Figure |
|---|---|---|---|---|
| FY3349 | MATa | his3Δ200 lys2-128δ ura3-52 leu2Δ0 hENT::AUR1C GDP-TK::URA3 SPT6-V5-AID::KanMX LEU2::TIR | This Study | Figure 1, S1 |
| FY3350 | MATa | his3Δ200 lys2-128δ ura3-52 leu2Δ0 MCM4-5xV5 bar1Δ::URA3 | This Study | Figure 3, 4 |
| FY3351 | MATa | his3Δ200 lys2-128δ ura3-52 leu2Δ0 SPT6-3xFLAG MCM4-5xV5 bar1Δ::URA3 | This Study | Figure 3, 4 |
| FY3352 | MATa | his3Δ200 lys2-128δ ura3-52 leu2Δ0 spt6YW-3xFLAG mcm4-5xV5 | This Study | Figure 4 |
| FY3353 | MATa | his3Δ200 lys2-128δ ura3-52 leu2Δ1 mcm2-5xV5 bar1::URA3 | This Study | Figure 3 |
| FY3354 | MATa | his3Δ200 lys2-128δ ura3-52 leu2Δ0 SPT6-3xFLAG Mcm2-5xV5 bar1Δ::URA3 | This Study | Figure 4 |
| FY3355 | MATa | his3Δ200 lys2-128δ ura3-52 leu2Δ0 Mcm4-5xV5 spt6-YW | This Study | Figure 4 |
| FY3356 | MATa | his3Δ200 lys2-128δ ura3-52 leu2Δ0 Mcm4-5xV5 spt6-50 | This Study | Figure 4 |
| FY3357 | MATa | his3Δ200 lys2-128δ ura3-52 leu2Δ0 3XFLAG-Spt6 Mcm4 | This Study | Figure 4 |
| FY3358 | MATa | his3Δ200 lys2-128δ ura3-52 leu2Δ0 3XFLAG-spt6-50 Mcm4-5xV5 | This Study | Figure 4 |
| FY3359 | MATa | his3Δ200 lys2-128δ ura3-52 leu2Δ0 Spt6-3xFLAG Mcm4-5xV5 rpb1::FSP | This Study | Figure 4 |
| FY3360 | MATa | his3Δ200 lys2-128δ ura3-52 leu2Δ0 Mcm4-5xV5 rpb1::FSP | This Study | Figure 4 |
| FY3361 | MATa | his3Δ200 lys2-128δ ura3-52 leu2Δ0 | This Study | Figure 1, 5, Supplemental Table 1 |
| FY3362 | MATa | his3Δ200 lys2-128δ ura3-52 leu2Δ0 spt6-YW | This Study | Figure 1, 5, Supplemental Table 1 |
| FY3363 | MATa | his3Δ200 lys2-128δ ura3-52 leu2Δ0 spt6-140 | This Study | Figure 1, Supplemental Table 1 |
| FY3364 | MATa | his3Δ200 lys2-128δ ura3-52 leu2Δ0 spt6-1004 | This Study | Figure 1, Supplemental Table 1 |
| FY3365 | MATa | his3Δ200 lys2-128δ ura3-52 leu2Δ0 spt6-50 | This Study | Figure 1, 5, Supplemental Table 1 |
| FY3371 | MATa | his3Δ200 lys2-128δ ura3-52 leu2Δ0 ctf4Δ::KanMX | This Study | Figure 5, Supplemental Table 1 |
| FY3372 | MATa | his3Δ200 lys2-128δ ura3-52 leu2Δ0 ctf4Δ::KanMX spt6-YW | This Study | Figure 5, Supplemental Table 1 |
| FY3373 | MATa | his3Δ200 lys2-128δ ura3-52 leu2Δ0 ctf4Δ::KanMX spt6-140 | This Study | Supplemental Table 1 |
| FY3374 | MATa | his3Δ200 lys2-128δ ura3-52 leu2Δ0 ctf4Δ::KanMX spt6-1004 | This Study | Supplemental Table 1 |
| FY3375 | MATa | his3Δ200 lys2-128δ ura3-52 leu2Δ0 ctf4Δ::KanMX spt6-50 | This Study | Figure 5, Supplemental Table 1 |
| FY3376 | MATα | his3Δ200 lys2-128δ ura3Δ0 leu2Δ0 mrc1Δ::KanMX | This Study | Figure 5, Supplemental Table 1 |
| FY3377 | MATα | his3Δ200 lys2-128δ ura3Δ0 leu2Δ0 mrc1Δ::KanMX spt6-YW | This Study | Figure 5, Supplemental Table 1 |
| FY3378 | MATα | his3Δ200 lys2-128δ ura3Δ0 leu2Δ0 mrc1Δ::KanMX spt6-140 | This Study | Supplemental Table 1 |
| FY3379 | MATa | his3Δ200 lys2-128δ ura3Δ0 leu2Δ0 mrc1Δ::KanMX spt6-1004 | This Study | Supplemental Table 1 |
| FY3380 | MATα | his3Δ200 lys2-128δ ura3Δ0 leu2Δ0 mrc1Δ::KanMX spt6-50 | This Study | Figure 5, Supplemental Table 1 |
| FY3381 | MATa | his3Δ200 lys2-128δ ura3-52 leu2Δ1 rad9Δ::URA3 trp1D63 | This Study | Figure 5, Supplemental Table 1 |
| FY3382 | MATa | his3Δ200 lys2-128δ ura3-52 leu2Δ1 rad9Δ::URA3 spt6-YW | This Study | Figure 5, Supplemental Table 1 |
| FY3383 | MATα | his3Δ200 lys2-128δ ura3-52 leu2Δ1 rad9Δ::URA3 spt6-140 | This Study | Supplemental Table 1 |
| FY3384 | MATa | his3Δ200 lys2-128δ ura3-52 leu2Δ1 rad9Δ::URA3 spt6-1004 | This Study | Supplemental Table 1 |
| FY3385 | MATa | his3Δ200 lys2-128δ ura3-52 leu2Δ1 rad9Δ::URA3 spt6-50 | This Study | Figure 5, Supplemental Table 1 |
| FY3386 | MATa | his3Δ200 lys2-128δ ura3-52 leu2Δ0 mad2Δ::KanMX | This Study | Supplemental Table 1 |
| FY3387 | MATα | his3Δ200 lys2-128δ ura3-52 leu2Δ0 mad2Δ::KanMX spt6-YW | This Study | Supplemental Table 1 |
| FY3388 | MATa | his3Δ200 lys2-128δ ura3-52 leu2Δ0 mad2Δ::KanMX spt6-140 | This Study | Supplemental Table 1 |
| FY3389 | MATa | his3Δ200 lys2-128δ ura3-52 leu2Δ0 mad2Δ::KanMX spt6-1004 | This Study | Supplemental Table 1 |
| FY3390 | MATa | his3Δ200 lys2-128δ ura3-52 leu2Δ0 mad2Δ::KanMX spt6-50 | This Study | Supplemental Table 1 |
| FY3392 | MATa | his3Δ200 lys2-128δ ura3-52 leu2Δ0 hENT::AUR1C GDP-TK::URA3 bar1Δ::KanMX | This Study | Fgure 2, S2 |
| FY3393 | MATa | his3Δ200 lys2-128δ ura3-52 leu2Δ0 hENT::AUR1C GDP-TK::URA3 spt6-YW bar1Δ::KanMX | This Study | Fgure 2, S2 |
| FY3394 | MATa | his3Δ200 lys2-128δ ura3-52 leu2Δ0 hENT::AUR1C GDP-TK::URA3 spt6-50 bar1Δ::KanMX | This Study | Fgure 2, S2 |
| FY3395 | MATα | his3::ITE::URA::LEU2 lys2-128δ ura3-52 leu2Δ0 | This Study | Figure 7, S3 |
| FY3396 | MATa | his3::ITE::URA::LEU2 lys2-128δ ura3-52 leu2Δ0 | This Study | Figure 7, S3 |
| FY3397 | MATa | his3::ITE::URA::LEU2 lys2-128δ ura3-52 leu2Δ0 spt6-YW | This Study | Figure 7, S3 |
| FY3398 | MATα | his3::ITE::URA::LEU2 lys2-128δ ura3-52 leu2Δ0 spt6-YW | This Study | Figure 7, S3 |
| FY3399 | MATa | his3::ITE::URA::LEU2 lys2-128δ ura3-52 leu2Δ0 spt6-140 | This Study | Figure 7, S3 |
| FY3400 | MATα | his3::ITE::URA::LEU2 lys2-128δ ura3-52 leu2Δ0 spt6-140 | This Study | Figure 7, S3 |
| FY3401 | MATα | his3::ITE::URA::LEU2 lys2-128δ ura3-52 leu2Δ0 spt6-1004 | This Study | Figure 7, S3 |
| FY3402 | MATa | his3::ITE::URA::LEU2 lys2-128d ura3-52 leu2D0 spt6-1004 | This Study | Figure 7, S3 |
| FY3403 | MATa | his3::ITE::URA::LEU2 lys2-128δ ura3-52 leu2Δ0 spt6-50 | This Study | Figure 7, S3 |
| FY3404 | MATα | his3::ITE::URA::LEU2 lys2-128δ ura3-52 leu2Δ0 spt6-50 | This Study | Figure 7, S3 |
| FY3405 | MATa | his3::INV::URA::LEU2 lys2-128δ ura3-52 leu2Δ0 | This Study | Figure 7, S3 |
| FY3406 | MATα | his3::INV::URA::LEU2 lys2-128δ ura3-52 leu2Δ0 | This Study | Figure 7, S3 |
| FY3407 | MATα | his3::INV::URA::LEU2 lys2-128δ ura3-52 leu2Δ0 spt6-YW | This Study | Figure 7, S3 |
| FY3408 | MATa | his3::INV::URA::LEU2 lys2-128δ ura3-52 leu2Δ0 spt6-YW | This Study | Figure 7, S3 |
| FY3409 | MATα | his3::INV::URA::LEU2 lys2-128δ ura3-52 leu2Δ0 spt6-140 | This Study | Figure 7, S3 |
| FY3410 | MATα | his3::INV::URA::LEU2 lys2-128δ ura3-52 leu2Δ0 spt6-140 | This Study | Figure 7, S3 |
| FY3411 | MATα | his3::INV::URA::LEU2 lys2-128δ ura3-52 leu2Δ0 spt6-1004 | This Study | Figure 7, S3 |
| FY3412 | MATα | his3::INV::URA::LEU2 lys2-128δ ura3-52 leu2Δ0 spt6-1004 | This Study | Figure 7, S3 |
| FY3413 | MATa | his3::INV::URA::LEU2 lys2-128δ ura3-52 leu2Δ0 spt6-50 | This Study | Figure 7, S3 |
| FY3414 | MATα | his3::INV::URA::LEU2 lys2-128δ ura3-52 leu2Δ0 spt6-50 | This Study | Figure 7, S3 |
| FY3416 | MATa | his3Δ200 lys2-128δ ura3Δ0 leu2Δ1 ade8-104 lig4Δ::KanMX | This Study | Supplemental Table 1 |
| FY3417 | MATa | his3Δ200 lys2-128δ ura3Δ0 leu2Δ1 ade8-104 lig4Δ::KanMX spt6-YW | This Study | Supplemental Table 1 |
| FY3418 | MATa | his3Δ200 lys2-128δ ura3Δ0 leu2Δ1 ade8-104 lig4Δ::KanMX spt6-140 | This Study | Supplemental Table 1 |
| FY3419 | MATa | his3Δ200 lys2-128δ ura3Δ0 leu2Δ1 ade8-104 lig4Δ::KanMX spt6-1004 | This Study | Supplemental Table 1 |
| FY3420 | MATa | his3Δ200 lys2-128δ ura3-52 leu2Δ1 ade8-104 lig4Δ::KanMX spt6-50 | This Study | Supplemental Table 1 |
| FY3421 | MATa | his3Δ200 lys2-128δ ura3-52 leu2Δ0 Rad52-eYFP::KanmX Htb1-cfp::Hyg | This Study | Figure 6 |
| FY3422 | MATα | his3Δ200 lys2-128δ ura3-52 leu2Δ0 Rad52-eYFP::KanmX Htb1-cfp::Hyg spt6-YW | This Study | Figure 6 |
| FY3423 | MATa | his3Δ200 lys2-128δ ura3-52 leu2Δ0 Rad52-eYFP::KanmX Htb1-cfp::Hyg spt6-1004 | This Study | Figure 6 |
| FY3424 | MATα | his3Δ200 lys2-128δ ura3-52 leu2Δ0 Rad52-eYFP::KanmX Htb1-cfp::Hyg spt6-50 | This Study | Figure 6 |
| FY3425 | MATa | his3Δ200 lys2-128δ ura3-52 leu2Δ0 Rad52-eYFP::KanmX Htb1-cfp::Hyg spt6-140 | This Study | Figure 6 |
| FY3426 | MATα | his3Δ200 lys2-128δ ura3-52 leu2Δ0 rad52Δ::KanMX | This Study | Figure 6, Supplemental Table 1 |
| FY3427 | MATa | his3Δ200 lys2-128δ ura3-52 leu2Δ0 rad52Δ::KanMX spt6-YW | This Study | Figure 6, Supplemental Table 1 |
| FY3428 | MATa | his3Δ200 lys2-128δ ura3-52 leu2Δ0 rad52Δ::KanMX spt6-140 | This Study | Supplemental Table 1 |
| FY3429 | MATa | his3Δ200 lys2-128δ ura3-52 leu2Δ0 rad52Δ::KanMX spt6-1004 | This Study | Supplemental Table 1 |
| FY3430 | MATa | his3Δ200 lys2-128δ ura3-52 leu2Δ0 rad52Δ::KanMX spt6-50 | This Study | Figure 6, Supplemental Table 1 |
| FY3431 | MATa | his3Δ200 lys2-128δ ura3-52 leu2Δ0 rnh1Δ::NatMX | This Study | Supplemental Table 1 |
| FY3432 | MATa | his3Δ200 lys2-128δ ura3-52 leu2Δ0 rnh201Δ::Hyg | This Study | Supplemental Table 1 |
| FY3433 | MATa | his3Δ200 lys2-128δ ura3-52 leu2Δ0 rnh1Δ::NatMX rnh201Δ::Hyg | This Study | Supplemental Table 1 |
| FY3434 | MATα | his3Δ200 lys2-128δ ura3-52 leu2Δ0 rnh1Δ::NatMX spt6-YW | This Study | Supplemental Table 1 |
| FY3435 | MATα | his3Δ200 lys2-128δ ura3-52 leu2Δ0 rnh201Δ::Hyg spt6-YW | This Study | Supplemental Table 1 |
| FY3436 | MATa | his3Δ200 lys2-128δ ura3-52 leu2Δ0 rnh1Δ::NatMX rnh201Δ::Hyg spt6-YW | This Study | Supplemental Table 1 |
| FY3437 | MATa | his3Δ200 lys2-128δ ura3-52 leu2Δ0 rnh1Δ::NatMX spt6-140 | This Study | Supplemental Table 1 |
| FY3438 | MATα | his3Δ200 lys2-128δ ura3-52 leu2Δ0 rnh201Δ::Hyg spt6-140 | This Study | Supplemental Table 1 |
| FY3439 | MATα | his3Δ200 lys2-128δ ura3-52 leu2Δ0 rnh1Δ::NatMX rnh201Δ::Hyg spt6-140 | This Study | Supplemental Table 1 |
| FY3440 | MATa | his3Δ200 lys2-128δ ura3-52 leu2Δ0 rnh1Δ::NatMX spt6-1004 | This Study | Supplemental Table 1 |
| FY3441 | MATα | his3Δ200 lys2-128δ ura3-52 leu2Δ0 rnh201Δ::Hyg spt6-1004 | This Study | Supplemental Table 1 |
| FY3442 | MATa | his3Δ200 lys2-128δ ura3-52 leu2Δ0 rnh1Δ::NatMX rnh201Δ::Hyg spt6-1004 | This Study | Supplemental Table 1 |
| FY3443 | MATa | his3Δ200 lys2-128δ ura3-52 leu2Δ0 rnh1Δ::NatMX spt6-50 | This Study | Supplemental Table 1 |
| FY3444 | MATa | his3Δ200 lys2-128δ ura3-52 leu2Δ0 rnh201Δ::Hyg spt6-50 | This Study | Supplemental Table 1 |
| FY3445 | MATα | his3Δ200 lys2-128δ ura3-52 leu2Δ0 rnh1Δ::NatMX rnh201Δ::Hyg spt6-50 | This Study | Supplemental Table 1 |
| FY3446 | MATα | his3::INV::URA::LEU lys2-128δ ura3-52 leu2Δ::ACT1p::Z3EV::Nat trp1Δ63 | This Study | Figure 7 |
| FY3447 | MATa | his3::INV::URA::LEU lys2-128δ ura3-52 leu2Δ::ACT1p::Z3EV::Nat trp1Δ63 spt6-YW | This Study | Figure 7 |
| FY3448 | MATα | his3::INV::URA::LEU lys2-128δ ura3-52 leu2Δ::ACT1p::Z3EV::Nat trp1Δ63 spt6-140 | This Study | Figure 7 |
| FY3449 | MATα | his3::INV::URA::LEU lys2-128δ ura3-52 leu2Δ::ACT1p::Z3EV::Nat trp1Δ63 spt6-1004 | This Study | Figure 7 |
| FY3450 | MATa | his3::INV::URA::LEU lys2-128δ ura3-52 leu2Δ::ACT1p::Z3EV::Nat trp1Δ63 spt6-50 | This Study | Figure 7 |
| FY57 | MATa | his4-192δ lys2-128δ ura3-52 | | Figure 7 |
| FY3207 | MATα | leu2Δ1 lys2-128δ ura3-52 spt6-YW | | Figure 7 |
| FWP10 | h- | wild type | | Figure 7 |
| yFS240 | h- | leu1-32 ura4-Δ18 ade6-210 his7-366 pJL218 (his7 adh1:tk) pFS181 (leu1 adh1:hENT1)) | Rhind lab | Figure 2 |

Supplmental Table 2. Related to STAR Methods. Oligonucleotides used for yeast strain construction.

| Strain Construction | | | | |
|---|---|---|---|---|
| Allele | DNA template | F | R | |
| *MCM4-5xV5* | pZM474 (Moqtaderi and Struhl, 2008 Yeast) | CGAGGGTGTAAGGAGATCAGTTCGCCTGAATAACCGTGTCCTGGAGCTCGGTAAACCTAT | ATTGTTACGCAGGGAATGATTGTAGTAGACAGCATCATCATAGGGCGAATTGGGTACCGG | |
| *MCM2-5xV5* | pZM474 (Moqtaderi and Struhl, 2008 Yeast) | CCAACTCCGCAGGTCTTTCGCAATTTATACCTTGGGTCACCTGGAGCTCGGTAAACCTAT | CATATCCAGATATTCGTAGGAATAACAAAGTTTTTCATCAAAGGGCGAATTGGGTACCGG | |
| *bar1Δ::URA3* | FY945 | AAAGTGAAGAGAAAGCACGT | CAAAATTGTGATGGCTGCAT | |
| *bar1Δ::KanMX* | pFA-KANMX | TTGATTAACGTGGATGAGTCCTTAAGAAGGCCGTTGAAGGGTCGACGGATCCCCGGGTT | CATACTAAATGGTGAAAGCGCGGAATCTTGGATCTAAACTCGATGAATTCGAGCTCGTT | |
| *rnh1Δ::natMX* | LW6838a (Koshland lab) | TGACAGCAGCATCAACAATG | AATCAACTGTGGACGAGGTT | |
| *rnh201Δ::hygMX* | LW6838a (Koshland lab) | CTTTGGCTGTGTGGATGATG | TCTTCTAGTCGTTCCGGTTG | |
| *ctf4Δ::KanMX* | pFA-KANMX | ATTGAGAAGGGCAAGAAGTGACGTAAATATACTAGACGTAGGTCGACGGATCCCCGGGTT | GGAGCATTTTGAACGATGATTTGAACAAATGAACAGGTATTCGATGAATTCGAGCTCGTT | |
| *mrc1Δ::KanMX* | pFA-KANMX | TCGTTATTCGCTTTTGAACTTATCACCAAATATTTTAGTGGGTCGACGGATCCCCGGGTT | AATGCGACTACTTCAAGACAGCTTCTGGAGTTCAATCAACTCGATGAATTCGAGCTCGTT | |
| *rad9Δ::URA3* | FY1944 | CTTTCTTATTACTGGCACGG | AAAGACCTTACCAACGTTGT | |
| *mad2Δ:KanMX* | pFA-KANMX | GTATTGAAAACCACTTCAAAGGGGCCCAATAGCACATTTAGGTCGACGGATCCCCGGGTT | TTGAATTCTATTAATATTTATAGCTGACCTGCGCACCAACTCGATGAATTCGAGCTCGTT | |
| *lig4Δ:KanMX* | Yeast ORF deletion set | CGGTTTCATCACTTACGTTG | ATAATTGCGCATCTTCCACT | |
| *rad52-eYFP-KanMX* | pYM39 (Janke et al 2004 Yeast) | AATAAATAATGATGCAAATTTTTTATTTGTTTCGGCCAGGAAGCGTTTCACCCATCGATGAATTCGAGCTCG | GAGAAGTTGGAAGACCAAAGATCAATCCCCTGCATGCACGCAAGCCTACTCGTACGCTGCAGGTCGAC | |
| *htb1-cfp-HygMX* | pYM30 (Janke et al 2004 Yeast) | GAAGGTACTAGAGCTGTTACCAAGTACTCTTCCTCTACTCAAGCAatcggtgacggtgctggtttaat | ATATACCCATATAAATAATAATATTAATTATAACCAAAGGAAGTGATTTCAggcggcgttagtatcgaatcgacag | |
| *rad52Δ::KanMX* | pFA-KANMX | AAGAACTGCTGAAGGTTCTGGTGGCTTTGGTGTGTTGTTGGAGCTCGTTTTCGACACTGG | AATGATGCAAATTTTTTATTTGTTTCGGCCAGGAAGCGTTTCCTTACCATTAAGTTGATC | |
| CRISPR-Cas9 mutagenesis alleles | sgRNA seq | Repair template F | Repair template R | Repair template GeneBlock |
| *rpb1-FSP* | GTTTTCACCTCTGGTTGATT | GGTGGCGTCACACCATACAGTAACGAAAGTGGTTTGGTCAATGCAGATCTTGACGTTAAA | ATTTGAACCTGAATCTTTTCTACGTCTCTTCATTTTCTCATCTTTAACGTCAAGATCTGC | |
| *3X-FLAG-SPT6 or 3X-FLAG-spt6-50* | TTTGCCTTTTATGGAAGAGA | | | GCAAACCAAATTGACAAGAAACTGGACAACGGTTAAAGCAAAAGGAGGAAGAAAGATATTGATTTGTGCTTTGAAGAATAACCCAAGCGAACAGGTCATTAATTGCCTAGAATCTTTACATAGAATTTGCCTTTTATGAGGGAACAAAAGCTGGAGCTCGACTATAAAGACGACGACGACAAAGCGGCGGATTACAAGGATGATGATGATAAGGCTGCAGATTATAAGGACGATGACGATAAGACTAGTCTCGAGGGGGGGCCCGGTACCCAATTCGCCCTAGAAGAGACGGGAGATTCGAAGCTGGTCCCTAGGGACGAGGAAGAAATAGTAAATGACAACGATGAAACTAAAGCGCCTAGTGAGGAAGAAGAAGGAGAAGATGTC |

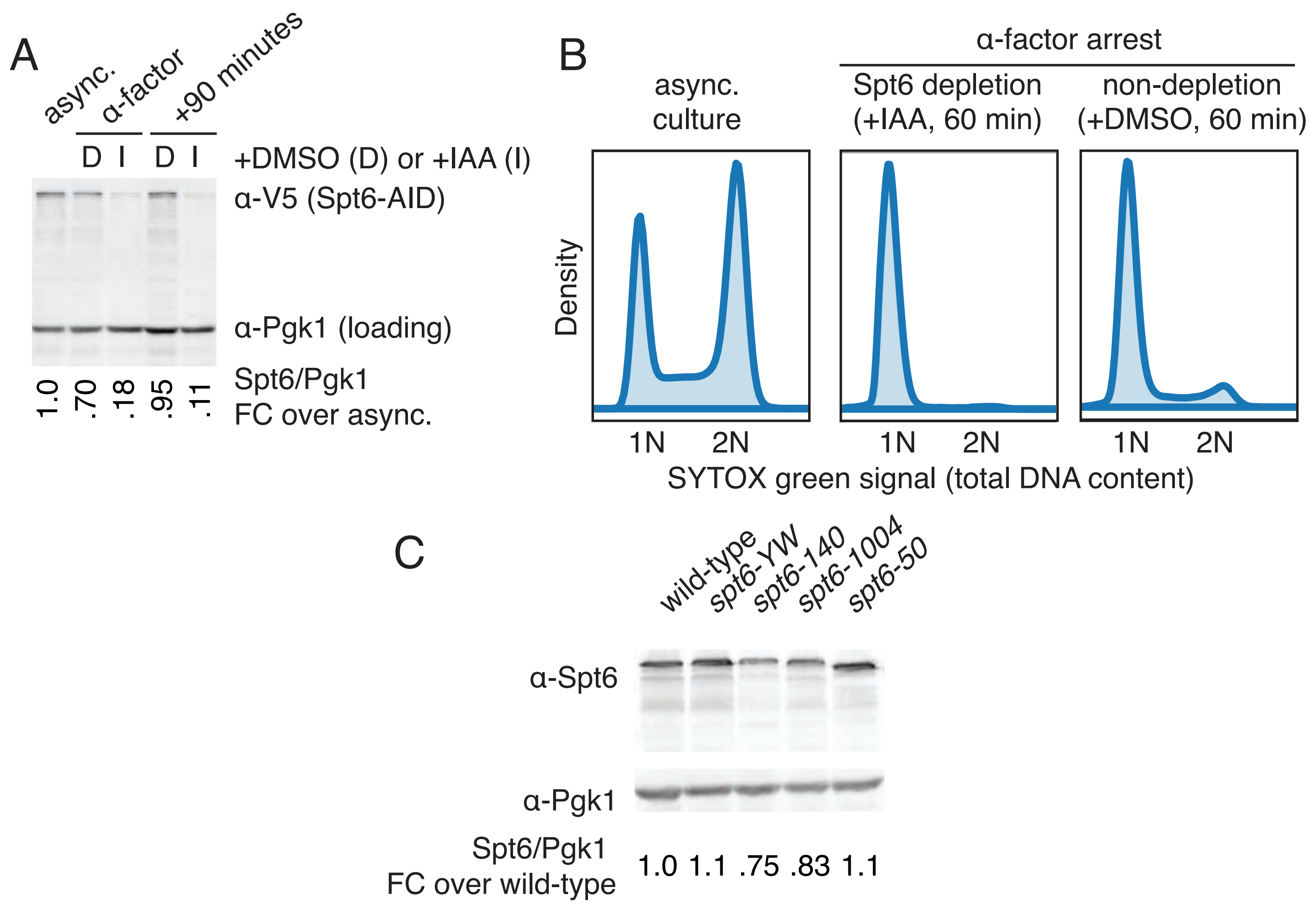

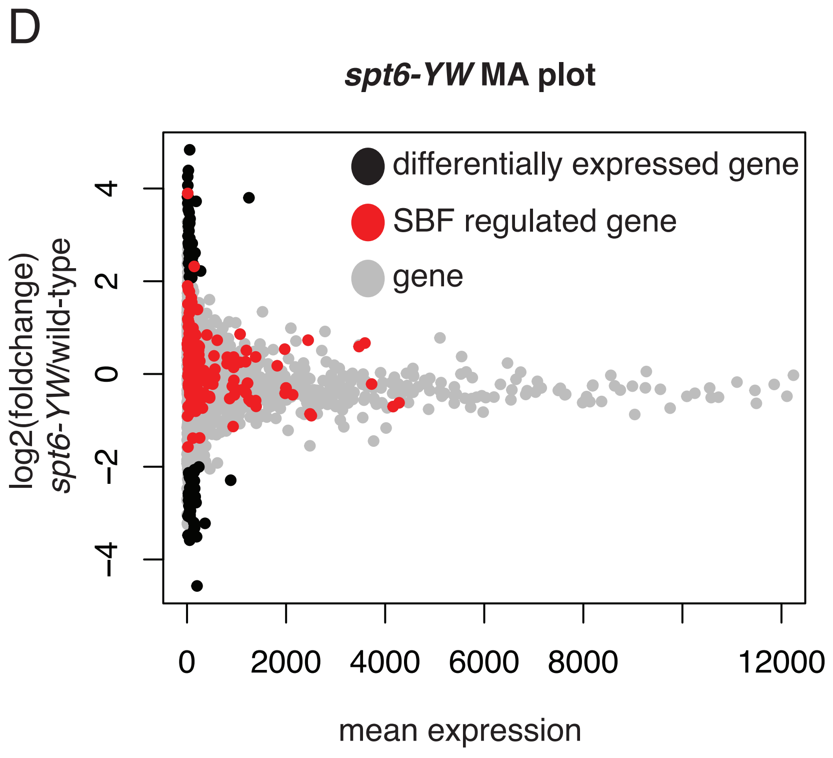

E

***spt6-YW* TSS-seq select genes**

| | log2FC | logpadj |
|---|---|---|
| *CLB1* | 0.777 | 0 |
| *CLB6* | 1.188 | 0 |
| *CLN1* | -0.685 | 0 |
| *CLN2* | -0.025 | 0 |
| *CLN3* | 0.855 | 0 |
| *RNR1* | 0.145 | 0 |
| *RNR2* | -0.136 | 0 |
| *RNR4* | -0.049 | 0 |
| *GFA1* | 3.893 | 1.596 |
| *PMA1* | 2.319 | 4.492 |

Supplemental Figure 1 Related to Figure 1 (A) A representative Western blot of Spt6-AID depletion. Spt6 levels are shown in asynchronously dividing samples (async.), after alpha-factor arrest in the presence of DMSO (D) or IAA (I) for one hour, and after release into fresh media for 90 minutes in the presence of DMSO or IAA. Pgk1 was used as a loading control. Quantification represents the average of biological triplicates. (B) Representative smoothed flow cytometry histogram for indicated time points and conditions. (C) A representative Western blot of the Spt6 protein levels for each spt6 mutant studied. Quantification of Spt6 protein levels in spt6 mutants. The average value of three biological replicates is shown. (D) TSS-seq data of spt6-YW vs. wild-type for strains grown at 30˚C (Viktorovskaya et al., 2021). The values for each gene are plotted based on their mean expression vs the log2(fold change) of spt6-YW / wild-type. Differentially expressed genes, defined as having a log2FC > 2, are indicated in black. Genes in red represent SBF targets (Iyer et al., 2001). (E) Shown are the log2 fold change (log2FC) and log adjusted p-values (logpadj) for select genes of interest – including cell cycle regulators, RNR genes, and the only SBF targets that are differentially expressed (GFA1, PMA1).

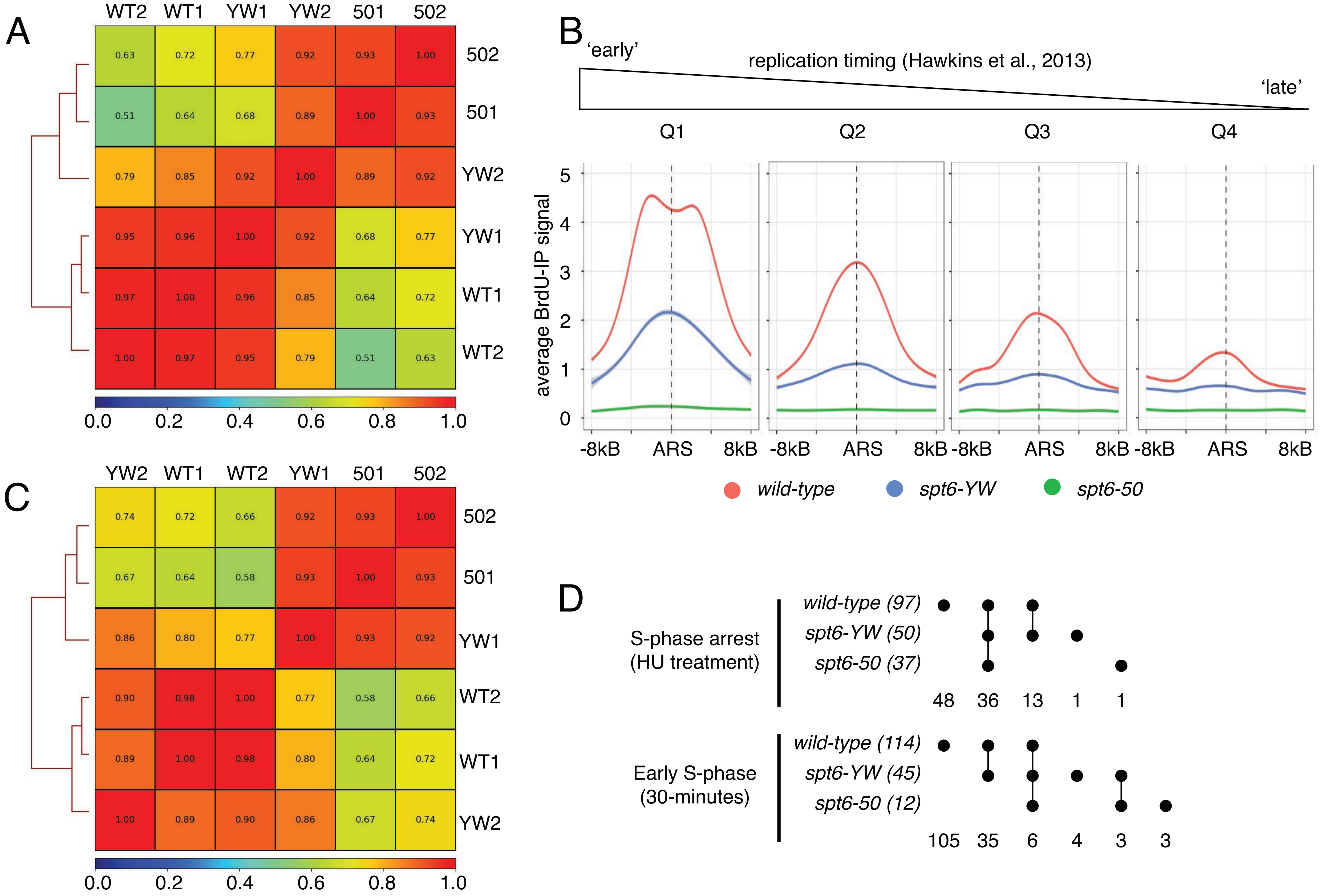

Supplemental Figure 2 Related to Figure 2 (A) Pearson correlation plots for individual BrdU-IP spike-in normalized IP signals for S-phase arrest (HU treatment) samples. (B) Replication origins were divided into quartiles based on previously reported replication timing (Hawkins et al., 2013), with an equal number of origins per group. The average value across all 58 ARSs per group is plotted for BrdU-IP S-phase arrested (HU treatment) samples (red for wild-type, blue for *spt6-YW*, and green for *spt6-50*). (C) Pearson correlation plots for individual BrdU-IP spike-in normalized IP signals for early S-phase (30 minute) samples. (D) Comparison of identified activated ARSs across genotypes. Genotypes are indicated on the left with the total number of activated ARSs in parenthesis. Samples are then compared in sets with the number of ARSs activated in that set indicated below the diagram.

| | permissive growth | elevated temperature | replication inhibitor | radiomimetic | alkylating agent | thymadine dimers |
|---|---|---|---|---|---|---|
| | YPD | YPD 37 | HU (100mM) | phleomycin (5 ug/mL) | MMS (0.024%) | UV (50 J/cm2) |
| wild-type | ++++ | ++++ | +++ | +++ | ++++ | ++++ |
| *spt6-YW* | ++++ | ++ | -/+ | ++ | + | ++++ |
| *spt6-140* | ++++ | -/+ | +++ | ++ | +++ | ++++ |
| *spt6-1004* | ++++ | +++ | +++ | ++ | +++ | ++++ |
| *spt6-50* | ++++ | ++++ | -/+ | - | - | ++++ |
| | YPD | YPD 37 | HU (10mM) | phleomycin (1 ug/mL) | MMS (0.003%) | UV (50 J/cm2) |
| *ctf4Δ* | ++++ | ++++ | ++++ | +++ | + | +++ |
| *spt6-YW ctf4Δ* | +++ | + | + | + | -/+ | +++ |
| *spt6-140 ctf4Δ* | ++++ | - | +++ | +++ | + | +++ |
| *spt6-1004 ctf4Δ* | ++++ | ++ | +++ | ++ | + | +++ |
| *spt6-50 ctf4Δ* | ++ | ++ | -/+ | - | - | +++ |
| | YPD | YPD 37 | HU (50mM) | phleomycin (1 ug/mL) | MMS (0.024%) | UV (50 J/cm2) |
| *mrc1Δ* | ++++ | ++++ | +++ | +++ | +++ | +++ |
| *spt6-YW mrc1Δ* | ++++ | - | + | ++ | + | +++ |
| *spt6-140 mrc1Δ* | ++++ | -/+ | +++ | +++ | +++ | +++ |
| *spt6-1004 mrc1Δ* | ++++ | +++ | ++ | +++ | +++ | +++ |
| *spt6-50 mrc1Δ* | +++ | ++ | -/+ | -/+ | - | +++ |
| | YPD | YPD 37 | HU (10mM) | phleomycin (1 ug/mL) | MMS (0.012%) | UV (50 J/cm2) |
| *rad9Δ* | ++++ | ++++ | ++++ | +++ | ++++ | +++ |
| *spt6-YW rad9Δ* | ++++ | ++ | ++ | ++ | +++ | +++ |
| *spt6-140 rad9Δ* | ++++ | -/+ | ++++ | ++ | +++ | +++ |
| *spt6-1004 rad9Δ* | ++++ | +++ | ++++ | ++ | ++ | +++ |
| *spt6-50 rad9Δ* | ++++ | ++++ | -/+ | - | + | -/+ |
| | YPD | YPD 37 | HU (100mM) | phleomycin (1 ug/mL) | MMS (0.024%) | UV (50 J/cm2) |
| *mad2Δ* | ++++ | ++++ | +++ | +++ | +++ | +++ |
| *spt6-YW mad2Δ* | ++++ | ++ | -/+ | +++ | ++ | +++ |
| *spt6-140 mad2Δ* | ++++ | -/+ | +++ | +++ | +++ | +++ |
| *spt6-1004 mad2Δ* | ++++ | +++ | +++ | +++ | +++ | +++ |
| *spt6-50 mad2Δ* | ++++ | ++++ | + | +++ | + | +++ |
| | YPD | YPD 37 | HU (10mM) | phleomycin (1 ug/mL) | MMS (0.003%) | UV (50 J/cm2) |
| *rad52Δ* | ++++ | ++++ | ++ | +++ | ++ | +++ |
| *spt6-YW rad52Δ* | +++ | - | -/+ | - | - | ++ |
| *spt6-140 rad52Δ* | ++++ | -/+ | ++ | +++ | ++ | +++ |
| *spt6-1004 rad52Δ* | ++++ | +++ | ++ | ++ | ++ | +++ |
| *spt6-50 rad52Δ* | ++ | +++ | -/+ | - | - | + |
| | YPD | YPD 37 | HU (100mM) | phleomycin (1 ug/mL) | MMS (0.024%) | UV (50 J/cm2) |
| *lig4Δ* | ++++ | ++++ | +++ | +++ | +++ | +++ |
| *spt6-YW lig4Δ* | ++++ | ++ | -/+ | +++ | ++ | +++ |
| *spt6-140 lig4Δ* | ++++ | -/+ | +++ | +++ | +++ | +++ |
| *spt6-1004 lig4Δ* | ++++ | +++ | +++ | +++ | +++ | +++ |
| *spt6-50 lig4Δ* | ++++ | ++++ | -/+ | +++ | ++ | +++ |
| | YPD | YPD 37 | HU (100mM) | phleomycin (1 ug/mL) | MMS (0.024%) | UV (50 J/cm2) |
| *rnh1Δ rnh201Δ* | ++++ | ++++ | ++ | +++ | + | +++ |
| *spt6-YW rnh1Δ rnh201Δ* | ++++ | ++ | -/+ | +++ | + | +++ |
| *spt6-140 rnh1Δ rnh201Δ* | ++++ | -/+ | ++ | +++ | + | +++ |
| *spt6-1004 rnh1Δ rnh201Δ* | ++++ | +++ | ++ | +++ | + | +++ |
| *spt6-50 rnh1Δ rnh201Δ* | ++++ | ++++ | - | + | -/+ | +++ |

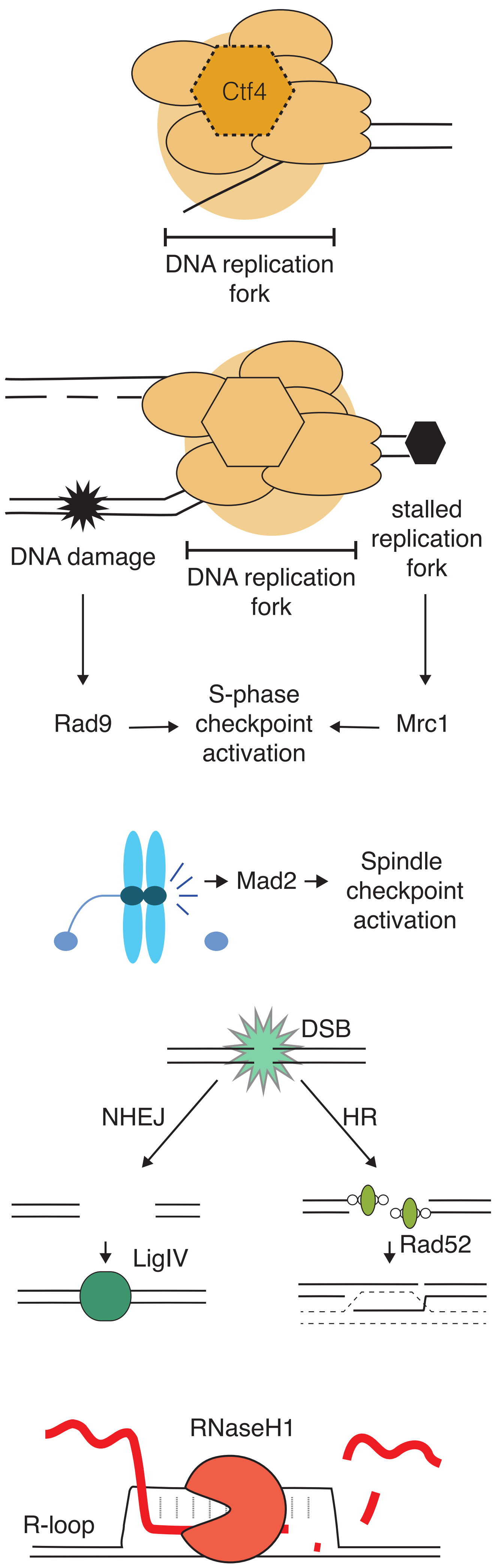

Supplemental Figure 3. Related to Figure 5. (Left) Summary of *spt6* genetic interactions. Indicated strains were spotted on indicated plates. Growth was summarized with + annotation for each visible circle complete growth. -/+ indicates partial growth and – indicates no detectable growth. Plates were scored on Day 2 or Day 3. (Right) Schemativ of genetic interactions tested.

A

| | with transcription (INV) | | | without transcription (ITE) | | |
|---|---|---|---|---|---|---|
| | average | stdev | FC over WT | average | stdev | FC over WT |
| *wild-type* | 8.9E-06 | 5.8E-06 | 1.0 | 3.9E-05 | 1.3E-05 | 1.0 |
| *spt6-YW* | 3.1E-05 | 2.3E-05 | 3.5 | 5.5E-05 | 1.5E-05 | 1.1 |
| *spt6-140* | 6.7E-05 | 2.1E-05 | 7.5 | 1.3E-04 | 8.0E-05 | 6.2 |
| *spt6-1004* | 3.2E-05 | 1.8E-05 | 3.6 | 6.7E-05 | 4.0E-05 | 3.1 |
| *wild-type* | 5.0E-05 | 1.1E-05 | 1.0 | 2.8E-05 | 1.1E-05 | 1.0 |
| *spt6-50* | 1.5E-03 | 6.0E-04 | 30.3 | 7.7E-04 | 1.0E-03 | 27.1 |

**B**

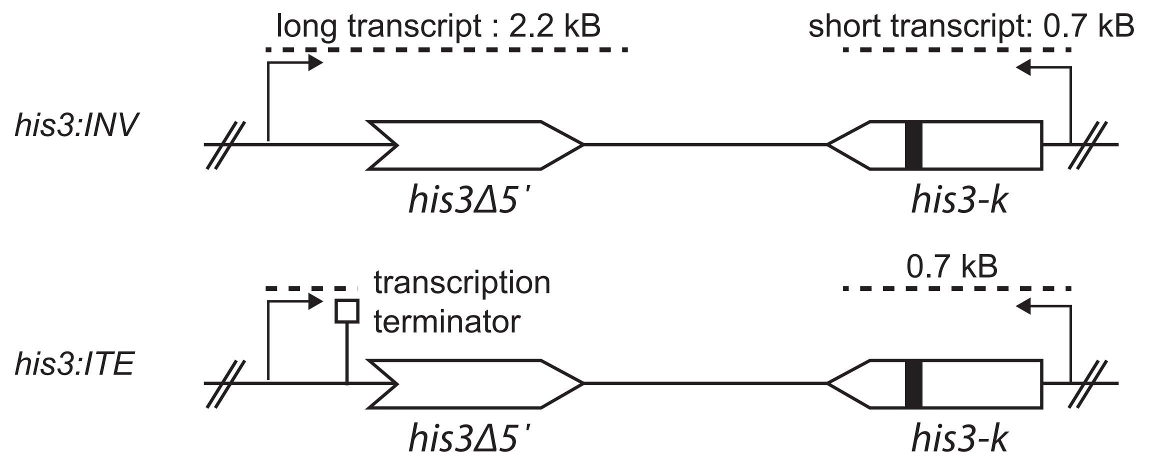

**C**

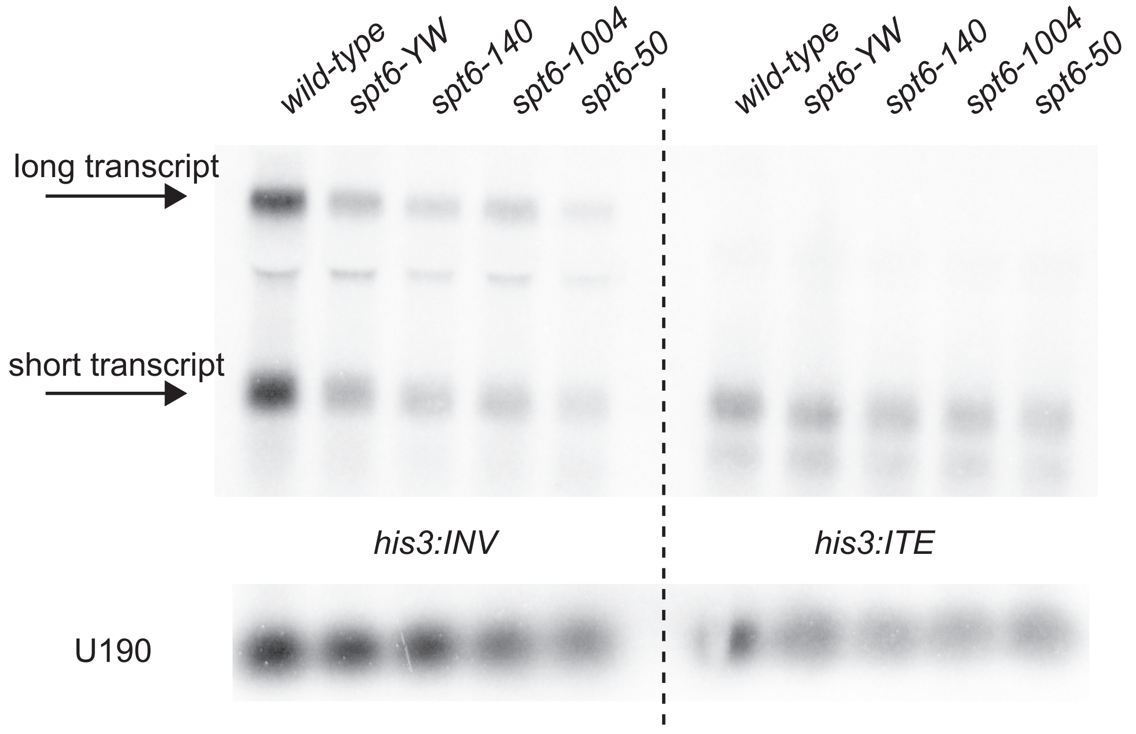

Supplemental Figure 4. Related to Figure 7. (A) Average values for all experiments in Figure 7A and 7B. (B) Schematic of reporters. (C) Representative Northern blot analysis of hyper-recombination reporter (Malagón and Aguilera, 2001).
